# VCAN is essential for ERK5-driven tumorigenesis in soft tissue sarcoma

**DOI:** 10.1101/2025.05.28.654281

**Authors:** Jaime Jiménez-Suárez, Francisco Cimas, José Joaquín Paricio, Borja Belandia, Yosra Berrouayel Dahour, Elena Arconada-Luque, Sofía Matilla-Almazán, Cesare Soffientini, Stefano Percio, Silvia Redondo-García, Natalia García-Flores, Cristina Garnés-García, Pablo Fernández-Aroca, Juan Jesús Martínez-Gómez, Antonio Fernández-Aramburo, Syong Hyun Nam-Cha, Elisabetta Rovida, Atanasio Pandiella, Azucena Esparís-Ogando, Sandro Pasquali, Juan Carlos Rodríguez-Manzaneque, Luis del Peso, María José Ruiz-Hidalgo, Ricardo Sánchez Prieto

## Abstract

The ERK5 signaling pathway has recently emerged as a critical regulator of soft tissue sarcoma (STS) biology, contributing to tumor initiation, progression, and maintenance. In this study, we identify VCAN, a chondroitin sulfate proteoglycan, as a novel transcriptional target of ERK5 and a central mediator of ERK5-related oncogenesis. Through a combination of genetic (silencing, overexpression) and pharmacological approaches, applied in both a chemically induced murine sarcoma model and several human STS cell lines, we demonstrate that ERK5 positively regulates VCAN expression. Functionally, VCAN silencing (by shRNAs) recapitulates the phenotypes of ERK5 silencing, including impaired migration, adhesion, proliferation, and tumorigenesis. Conversely, VCAN overexpression rescues these effects, confirming its essential role in ERK5-mediated oncogenesis. Furthermore, transcriptomic profiling reveals that VCAN accounts for a substantial portion of ERK5-regulated gene expression program. Analyses of human STS patient samples reveal significantly elevated mRNA levels of both VCAN and ERK5 compared to normal tissues. Notably, a strong correlation between VCAN and ERK5 expression, both at mRNA and protein levels, emerged in biopsies from leiomyosarcomas and undifferentiated pleomorphic sarcomas. Together, these findings uncover VCAN as a key effector in ERK5-driven tumorigenesis and highlight the ERK5/VCAN signaling axis as a promising therapeutic target in soft tissue sarcomas.

## Introduction

Soft tissue sarcomas (STS) represent a diverse group of tumors originating from the embryonic mesodermal, accounting for approximately 1% of all adult solid malignant cancers and about 15% of all pediatric tumors (1). Despite intense research aimed at improving the outcome of the disease, therapy of STS still achieves poor results, highlighting the need for better knowledge of the pathophysiological entities that contribute to its progression. Identification of the molecular alterations that drive STS progression may facilitate the development of targeted strategies that could improve patient prognosis. While certain types of STS such as Ewing’s sarcoma are well-characterized (2), the molecular bases of STS are not fully understood, probably due to their high heterogeneity. Therefore, ongoing research efforts aim to elucidate the molecular bases of these tumors, to better classify the various STS histologies (3).

Beyond genetic factors, cellular signaling alterations have emerged as key contributors to sarcoma biology. In this regard, the mitogen-activated protein kinases (MAPKs) family, one of the main nodes in cellular signaling which includes ERK1/2, P38, JNK and ERK5, has been extensively studied in the context of STS biology (4–7). Regarding the latter, recent studies have shown a determinant role of ERK5 signaling in STS by *in vivo* carcinogenesis, using 3Methyl-cholantrene (3MC), and a genetically modified mouse model with a constitutively active ERK5 signaling pathway (8,9). In these experimental models, chemical carcinogenesis induced pleomorphic sarcomas with muscle differentiation, histologically resembling human leiomyosarcoma (LMS). On the other hand, transgenic mice expressing constitutively active MEK5, the ERK5 upstream activating kinase, developed undifferentiated pleomorphic sarcomas (UPS). This evidence suggests that the ERK5 pathway may contribute, at least in some histologies of STS, to sarcomagenesis. However, the precise role of ERK5 in the variety of STS and its downstream targets are not fully clarified. In this regard, it has been shown that ERK5 depletion affects multiple biological processes (e.g. proliferation, motility, adhesion etc.) linked to the oncogenic phenotype. Indeed, a deeper understanding of ERK5-regulated effectors could contribute to the assessment of an ERK5-based therapy in sarcoma pathology. Interestingly, in primary cell cultures derived from the 3MC murine sarcoma model, VCAN was one of the significantly differentially expressed genes (DEG) after ERK5 silencing (8).

*VCAN* codes for Versican, a chondroitin sulfate/dermatan sulfate proteoglycan found in the extracellular matrix (ECM) and interstitial space of most tissues playing a critical role in key cellular processes such as proliferation, migration, adhesion, inflammation and immunity (10), all of which have been shown to be affected by ERK5 in several tumor types (reviewed in (11)). In humans, the *VCAN* gene encodes multiple isoforms (V0, V1, V2, V3 and V4), due to alternative splicing of exons 7 and 8 (12). Besides, the V1 isoform can be proteolyzed by ADAMTS proteases, generating an N-terminal fragment (versikine) with a plethora of additional biological functions (13). VCAN has been implicated in inflammatory disorders (14,15), vascular diseases (16,17) and certain genetic conditions like Wagner syndrome (18).

In this background, we evaluated the role of VCAN in mediating ERK5-associated biological processes such as proliferation, migration, adhesion and tumorigenesis *in vivo* in different experimental models of STS. Our data demonstrate that ERK5 regulates, at the transcriptional level, the expression of VCAN which is critical for the oncogenic characteristics of STS, as evidenced by cell culture and human samples studies. These findings highlight the signaling axis ERK5-VCAN as a potential therapeutic target, offering new opportunities for intervention in STS.

## Materials and methods

### Cell Lines

Human HEK-293T, SK-LMS-1 (LhMS), 786-O (renal cell carcinoma) and Hs 578T (breast cancer) cell lines were purchased from ATCC (LGC, Barcelona, Spain). Cells were maintained in 5% CO_2_ at 37°C and grown in DMEM supplemented with 10% FBS, 1% glutamine plus antibiotics. AA and EC cell lines derived from LMS and rhabdomyosarcoma, respectively, were kindly provided by Dr. Carnero (IBIS, Sevilla, Spain). They were maintained in 5% CO_2_ at 37°C and grown in Ham’s Nutrient Mixture F10 supplemented with 10% FBS, 1% glutamine plus antibiotics. The murine sarcoma 3MC-C1 cell line has been previously described (8). Cell culture reagents were provided by Lonza (Cultek, Madrid, Spain).

### Plasmids, antibodies and chemicals

Plasmids used were as follows. For luciferase assay, pCEFL HA-ERK5-WT, pCEFL MEK5DD and PCEFL HA-ERK5-KD have been previously described (19), and pLightSwitch VCAN Promoter Reporter (Ref. S712930; Active Motif, Carlsbad, CA, USA). For shRNA assays, all plasmids, with a PLKO.1 basis, were purchased from Merck (Tres Cantos, Madrid, Spain): shRNA ERK5-1 Human (TRCN0000010275), shRNA ERK5-2 Human (TRCN0000197264), shRNA VCAN-1 Human (TRCN0000033637), shRNA VCAN-2 Human (TRCN0000033638), and for mouse cell lines shRNA ERK5 (TRCN0000232396), and shRNA VCAN (TRCN0000175477). For stable MEK5DD expression in SK-LMS-1 cells, the Flag-MEK5DD construct described in (20) was subcloned into a pBabe-puro vector. The plasmid for VCAN-V1 overexpression was kindly provided by Dr Dieter R. Zimmermann (21). Antibodies used are listed in Supplementary Table 1. Crystal violet was purchased from Merck. XMD8-92 and JWG-071 were obtained from Selleckchem (Deltaclon, Madrid, Spain). Puromycin was purchased from Merck and Zeocin was obtained from Thermo Fisher Scientific (Madrid Spain). Collagen was purchased from Advanced Biomatrix (Carlsbad, CA USA). Mimosine was purchased from MedChemExpress (Eurodiagnostico, Madrid, Spain).

### Animal Studies

All the animal experimentation was carried out according to Spanish (RD 53/2013) and European Union regulations (2010/63/UE) and approved by the Ethics in Animal Care Committee of the University of Castilla-La Mancha (reference ES020030000490). For xenograft assays, 5 × 10^5^ cells from 3MC-C1 or derived cell lines were subcutaneously injected into the back of 5/6-week-old female mice of the J:NMRIFoxn1nu/Foxn1nu strain (Janvier, France). In the case of SK-LMS-1 or derived cell lines, 2 × 10^6^ cells were subcutaneously injected into the back of 5/6-week-old NOD.Cg-Prkdc^SCI^ Il2rg^tm1Wjl^/SzJ female mice (Charles River, France). Tumors were measured by caliper twice a week, and tumor volume was calculated according to the formula V = (D × d^2^)/2 (where D is tumor length and d tumor width).

### Lentiviral and Retroviral Production and Infections

Lentiviral production and infections were performed as previously described (22). One day after infection, cells were selected with puromycin for 72 hours (SK-LMS-1: 1 µg/mL; AA: 1.5 µg/mL; EC: 1.9 µg/mL; 3MC-C1: 2.8 µg/mL; 786-O: 3 µg/mL; Hs578T: 1.3 µg/mL). Each experiment was performed with at least two different pools of infection. Infected cells were discarded 15 days after selection, and new pools were generated.

For retroviral production (pBabe constructs), HEK-293T cells were transfected using the jetPEI transfection reagent following provider’s instructions (Polyplus Transfection, Sélestat, France). To generate viruses, VSV, Gag-pool and the plasmid of interest (pBabe-MEK5DD or its control pBabe-puro) were used as in the case of shRNA. After 48 hours, target cells were infected and 72 hours later were selected by puromycin treatment before use.

### VCAN-V1 Overexpression

FuGENE HD Transfection Reagent, obtained from Promega, was used following manufacturer instructions. SK-LMS-1 cells were selected with 300 µg/mL of zeocin for a period of 4 weeks.

### Western Blotting

For VCAN analyses, media was collected from different cell lines after 48 hours in the absence of serum, and secreted proteins were concentrated with StrataClean Resin (400714, Agilent Technologies). Concentrated proteins were incubated 1 hour at 37°C with Chondroitinase ABC (C3667, Sigma-Aldrich), in Chondroitinase buffer (180 mM Tris, 216 mM Sodium Acetate) with Trypsin inhibitor (from chicken egg white, T9253, Sigma-Aldrich). After treatment, Laemmly buffer with β-mercaptoethanol was added and samples were heated for 10 minutes at 100°C. Finally, proteins were resolved in 4-20% Mini-Protean TGX Precast protein gels (BioRad) and transferred to Polyvinylidene difluoride (PVDF) membranes (BioRad). Membranes were blocked with 5% low-fat milk and incubated overnight with the indicated antibodies. After incubation with the appropriate secondary peroxidase-conjugated antibody, signal was detected with the Amersham ECL Prime Western Blotting Detection Reagent (GE Healthcare Life Sciences) in an ImageQuant LAS4000 (GE Healthcare Life Sciences). The rest of protein quantifications and western blots were performed as previously described (19).

### Clonogenic Assay

Clonogenic assays were performed by using 200-600 cells/well seeded in 6-well plates and maintained for 10–14 days. The colonies with less than 5 mm diameter or 50 cells were discarded.

### Growth Curves

Growth curves were performed as previously described (8). Briefly, 3 × 10^5^ cells for 3MC-C1 and SK-LMS-1, 2.5 × 10^5^ cells for AA and 7.5 × 10^5^ cells for EC were seeded into 100 mm plates and counted on days 3, 6, and 9 by using an automated cell counter (Bio-Rad) and replated in the same manner up to day 9. This experiment was performed with 3 different pools of infection for each cell line. Graphics show the cumulative cell number from a representative experiment out of 3 with nearly identical results in different pools of infections.

### Adhesion Assay

For adhesion assays, 24-well plates were coated with collagen at a concentration of 100 µg/mL per well, diluted in Milli-Q H_2_O, and incubated for 2 hours at room temperature. The collagen was then removed, and wells were washed twice with DPBS. Subsequently, 1.5 × 10^4^ SK-LMS-1 cells/well were seeded, and then imaged every 15 minutes using the Axio Observer microscope (Zeiss; Madrid Spain). Images were analyzed with the ImageJ software plug-in CellCounter, counting both cells attached and expanded and those that were not. Triplicates were performed for each condition and 3 independent experiments were developed.

### Luciferase Reporter Assays

SK-LMS-1 and HEK-293T cells were transfected with FuGENE HD (Promega) according to the manufacturer’s instructions. For the transient transcriptional assays, 1.2 × 10^4^ SK-LMS-1 cells and 4 × 10^4^ HEK-293T cells per well were seeded in 24-well plates 24 hours prior to transfect with 120 ng of the Renilla-luciferase reporter (pLightSwitch VCAN), 100 ng of pCEFL MEK5DD and/or 100 ng of pCEFL HA-ERK5-WT or KD. The amount of total DNA transfected at all points was matched with pCEFL empty plasmid. Twenty-four hours after transfection, the Renilla-luciferase activities were determined using the LightSwitch Luciferase Assay Kit (SwitchGear Genomics, Menlo Park, CA, USA) in a GLoMAX® luminometer (Promega) according to the manufacturer’s instructions.

### Migration Assay

Migration was evaluated by wound healing assay. SK-LMS-1 and derived cell lines were seeded in 24-well plates at 1.2 × 10^5^ cells/well. After 24 hours, a wound was carefully made in each well using a 10 µL tip. Culture medium was then removed, the wells were washed twice with DPBS, and 1 mL of DMEM containing 0.5% FBS was added. Wells were then imaged every hour using the Axio Observer microscope (Zeiss). The images were analyzed with the MRI WoundHealing Tool plug-in of ImageJ software. A triplicate was performed for each condition and 3 independent experiments were performed to obtain the result.

### Quantitative PCR (RT-qPCR)

Total RNA from cells and mice tumor samples (after tissue homogenization with a Polytron) was obtained as previously described (8). cDNA synthesis and PCR conditions were performed as indicated (8). For RT-qPCR, murine *B2m* and human *GAPDH* were used as endogenous controls. Primers were designed by using the NCBI BLAST software and purchased from Sigma-Aldrich. The primers used are listed in Supplementary Table 2.

### Human Samples, Histology and Immunohistochemistry

Patients’ cohort from Fondazione IRCCS Istituto Nazionale dei Tumori (INT) in Milan, Italy, comprised patients with STS enrolled in the retrospective arm of the SARCOMICS study. Initiated in 2018 at INT, SARCOMICS is an observational study designed to assess whether integrating radiomic, genomic, and immunological markers can improve the predictive accuracy of clinical-based nomograms. The study includes both retrospective and prospective cohorts of patients diagnosed with primary retroperitoneal sarcomas or primary extremity/superficial trunk STS who underwent curative-intent surgery. The retrospective cohort has been here exploited for analyses of this study. The study has been approved by the Institutional Ethics Committee at Fondazione IRCCS Istituto Nazionale Tumori, Milan, Italy (ID: INT 77/18).

For immunohistochemistry, human tumor samples were provided by the Tumor Bank of the Complejo Hospitalario Universitario de Albacete with the corresponding ethical committee approval (number 2021-131). The selection of cases was performed by retrospective search in the archive of the Pathology Department of the Complejo Hospitalario Universitario de Albacete, selecting 10 cases of LMS and 9 cases of UPS diagnosed between 2018 and 2023. For VCAN immunohistochemistry, deparaffinization and antigenic recovery were done using the pT-Link device, at 95°C for 20 minutes, after which staining was performed using the Autostainer Link 48 system, with a 1/2000 concentration and incubation period, using a linker for the secondary antibody. ERK5 immunohistochemistry was performed as previously described (8). Cases were classified as positive (+) or negative (0) if the percentage of tumor cells with protein expression was less than 10%. In positive cases, if staining was incomplete and/or weakly positive it was quantified as +, and in cases with complete expression it was reported as ++ and +++ if the intensity of labelling was moderate or intense, respectively. All cases were evaluated by two trained pathologists.

### RNA Sequencing (RNA-seq) and Transcriptomic Analysis

SK-LMS-1 cells were infected with PLKO.1-empty vector or PLKO.1-shRNA ERK5-1 or shRNA VCAN-2. Three different pools of infection were used. Total RNA was extracted as previously described and RNA integrity was determined by Agilent 2100 Bioanalyzer (RIN range, 9.1–9.9). Reverse stranded library preparation and RNA sequencing were conducted by BGI company using the DNBSEQ platform, generating paired-end 100-bp reads. Raw sequencing data were processed by BGI with SOAPnuke (version 1.5.2) (23) with the following parameters: -l 15, -q 0.2, -n 0.05 (https://github.com/BGI-flexlab/SOAPnuke). The processing steps included: 1) removal of reads containing adaptor sequences; 2) removal of reads with N content greater than 5%; and 3) removal of low-quality reads, defined as those with more than 20% of bases having a quality score below 15. Filtered “clean reads” were saved in FASTQ format.

The filtered reads were aligned to the human reference genome (GRCh38) using HISAT2 (version 2.1.0) (24) in paired-end mode with default settings. SAM files generated during alignment were converted to BAM format and sorted by coordinates using SAMtools (version 1.6) (25). Gene-level quantification was performed with HTSeq-count (version 0.11.3) (26) with the GRCh38.109 gene annotation in GTF format, specifying the reverse-stranded option.

Lowly expressed genes were filtered using the filterByExpr function from the edgeR package in R (version 4.3.3) (27) with default settings. Differential expression analysis was performed using the Limma-voom pipeline (28), and *p*-values were adjusted for multiple testing using the Benjamini-Hochberg method. Genes with an adjusted *p*-value < 0.05 were considered differentially expressed.

### Functional Enrichment Analysis

Gene Ontology (GO) enrichment analysis for Biological Processes was performed to assess functional enrichment of DEGs (adjusted *p*-value < 0.05) using clusterProfiler package in R (version 4.8.3) (29).

### Statistical Analysis

The data are reported as the mean ± standard deviation (SD) or the standard error of the mean (SEM). Statistical analysis was performed using GraphPadPrism 9 and Office Excel 2020 (Microsoft). Significance was determined using a t-test or non-parametric tests. The statistical significance of differences is indicated in Figures by asterisks as follows: * p < 0.05; ** p < 0.01; and *** p < 0.001. Correlation analysis was computed with Spearman coefficient and p values are indicated in figures.

## Results

### ERK5 regulates VCAN expression in sarcoma derived cell lines

We previously reported that in sarcoma cells derived from a 3MC murine sarcoma model, upon ERK5 silencing, the key ECM proteoglycan VCAN was among the significantly down-regulated differentially expressed genes (DEG) suggesting its contribution to the biological effects driven by ERK5 signaling (8). To investigate further, we used shRNA to knockdown *MAPK7* (encoding ERK5) or *VCAN* expression in our 3MC-C1 murine cell line with knockdown efficiency confirmed at both RNA and protein levels (Fig.1A and B). 3MC-C1 cells with diminished ERK5 expression showed the expected decrease in VCAN expression at mRNA and protein levels (Fig.1 B). Interestingly, *VCAN-*silenced cells phenocopied ERK5 abrogation in terms of reduced proliferation (Fig. 1C), diminished colony formation ability (Fig. 1D) and impaired *in vivo* tumorigenesis (Fig. 1E), suggesting a clear correlation between these two proteins at the biological level. To gain further insights into this putative connection, we switched to a human experimental model using the well-established SK-LMS-1 LMS cell line. In these cells, using two different shRNAs against *MAPK7*, we achieved efficient ERK5 knockdown at both RNA and protein levels (Fig. 2A). Again, similarly with what observed in the 3MC-C1 murine cell line, ERK5 knockdown correlated with a decreased *VCAN* mRNA level (Fig. 2B). Next, we decided to challenge the use of known ERK5 chemical inhibitors such as XMD8-92 and JWG-071 (for a review see (30)) observing a specific decrease in *VCAN* mRNA levels (Fig. 2C), as confirmed by the expected increase in *CDKN1A* mRNA (31). In fact, similar results were obtained in other cellular models of LMS (AA) and rhabdomyosarcoma (EC) (Supplementary Fig. 1). Furthermore, regulation of *VCAN* by ERK5 was confirmed in other cell lines derived from unrelated pathologies with epithelial origin such as breast (Hs 578T) or renal (786-O) cancers, yielding comparable results (Supplementary Fig. 2) supporting the broader relevance of our observation.

**Figure 1.**
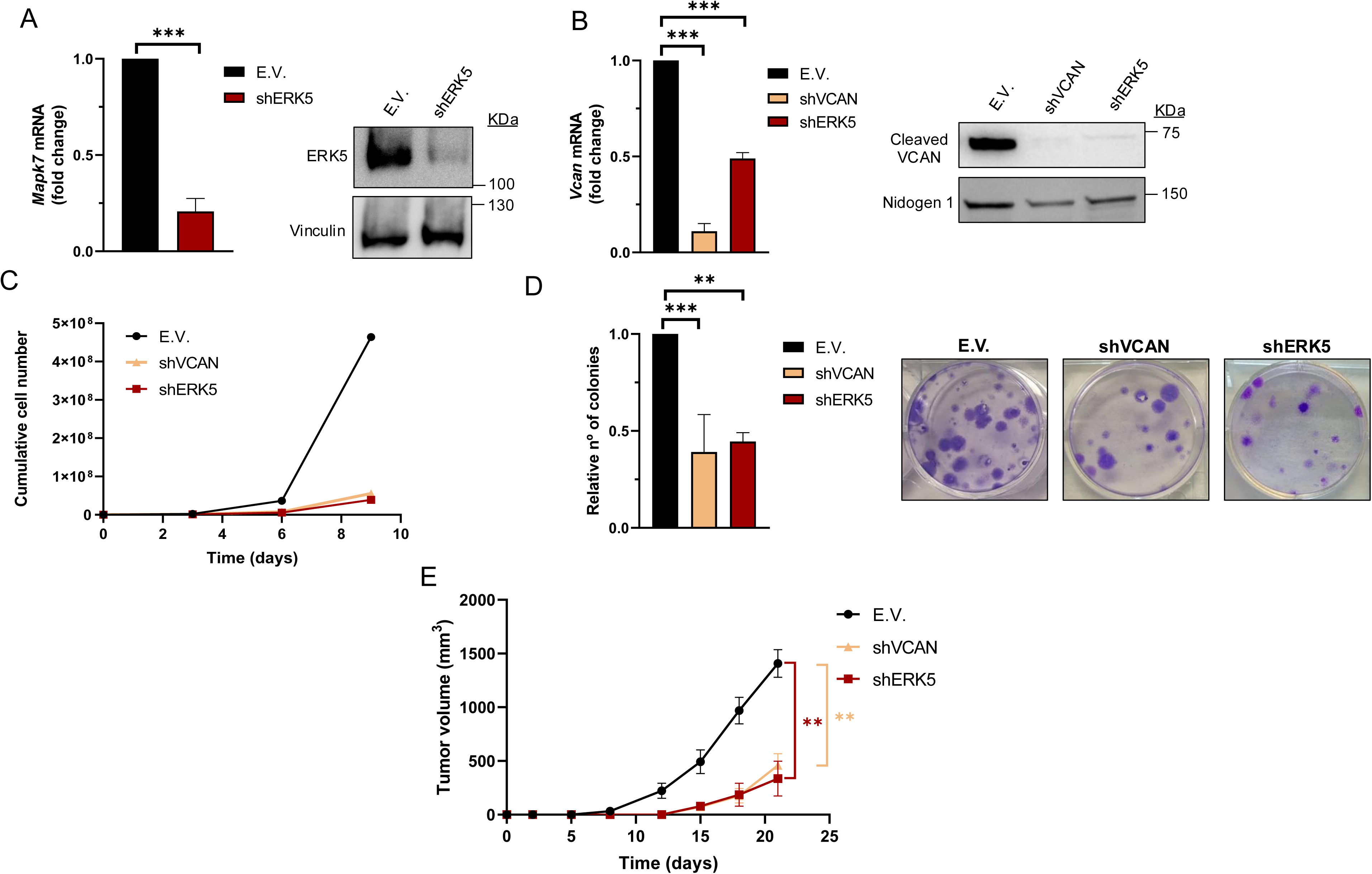
VCAN silencing mimics the *in vitro* and *in vivo* effects of ERK5 knockdown in the murine cell line 3MC-C1. 3MC-C1 cells were infected with lentiviruses carrying PLKO.1-empty vector (E.V.), the murine PLKO.1-shRNA ERK5 (shERK5) or PLKO.1-shVCAN (shVCAN). (A) ERK5 silencing relative mRNA levels were evaluated by RTqPCR (left panel) and protein levels by western blot (right panel), using Vinculin as a loading control. (B) VCAN relative mRNA (left panel) and protein levels (right panel) were analyzed by RT-qPCR and western blot, using Nidogen 1 as loading control. (C) Cumulative cell number experiment for 3 × 10^5^ E.V., shERK5 or shVCAN 3MC-C1 cells seeded in 100 mm plates and replated every 3 days up to day 9. A representative experiment out of 3 different pools of infections with nearly identical results is shown. (D) Clonogenic assay of E.V., shERK5 and shVCAN 3MC-C1 cells was performed by seeding 200 cells/well in a 6 well plate and stained with crystal violet after 12 days. Histogram shows the relative number of colonies obtained in clonogenic assays indicating the mean +/-SD of 3 independent pools of infection (left panel). Representative images of clonogenic assays for each condition are shown in right panel. (E) Tumor growth of nude mice (n=3) inoculated with 5 × 10^5^ cells of each cell line derived from 3MC-C1, measured at the indicated times. The graphic represents the mean +/-SEM. The unpaired Student’s t-test was used to assess statistical significance. **p<0.01; ***p<0.001.

**Figure 2.**
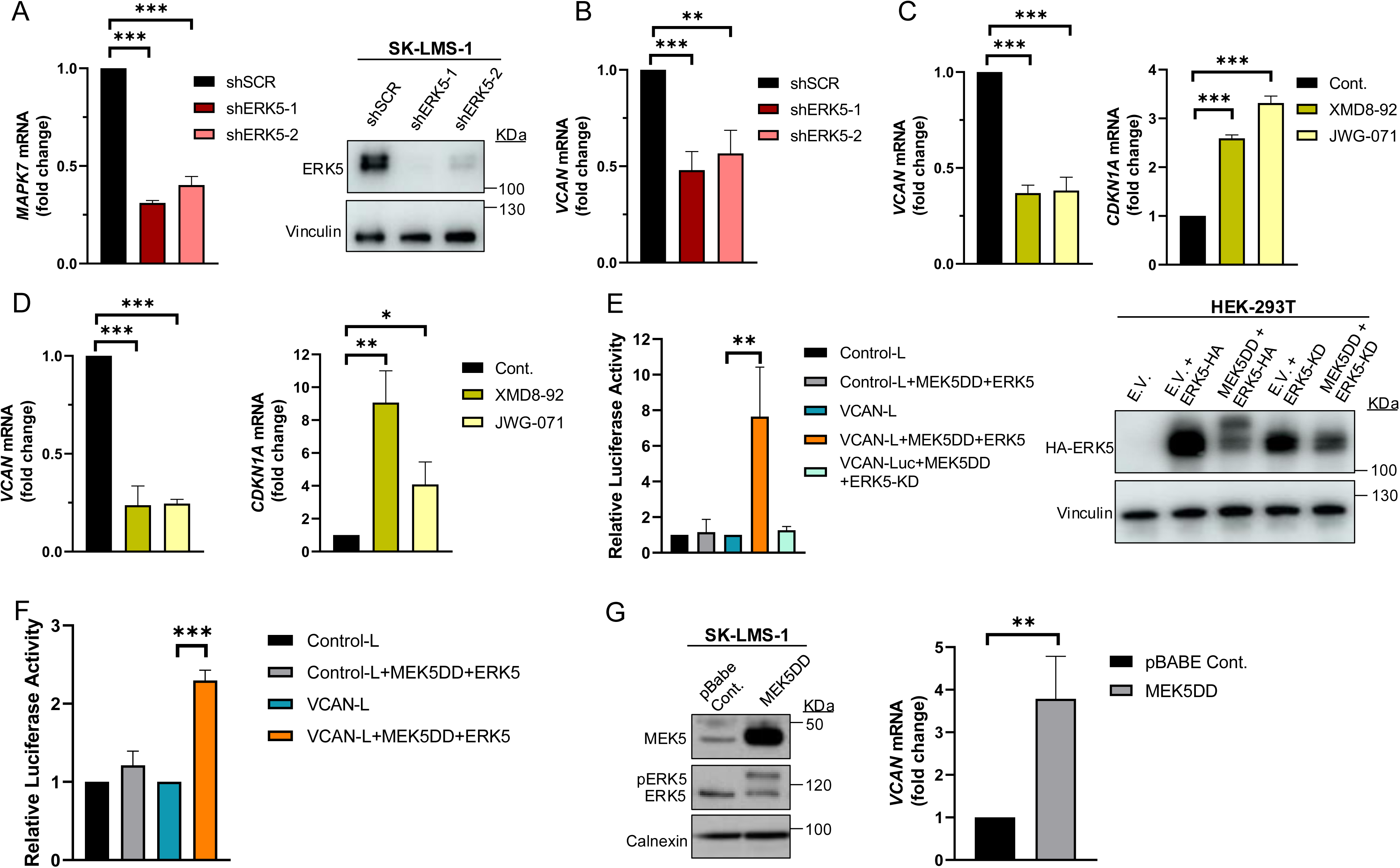
ERK5 regulates *VCAN* promoter activity in SK-LMS-1 cells. (A) Assessment of *MAPK7* interference in SK-LMS-1 cells by lentiviral infection carrying the PLKO.1-shScramble (shSCR) or shRNA for *MAPK7* (PLKO.1-shRNA ERK5-1/2) vectors. Relative mRNA levels were evaluated by RT-qPCR (left panel) and protein levels were detected by western blot using Vinculin as a loading control (right panel). (B) *VCAN* relative mRNA levels in SK-LMS-1 cells infected with shSCR and shERK5-1/2 measured by RT-qPCR. (C) *VCAN* (left panel) and *CDKN1A (*right panel) relative mRNA levels measured by RT-qPCR in SK-LMS-1 cells treated for 18 hours with XMD8-92 (5 µM) and JWG-071 (5 µM) inhibitors. (D) *VCAN* (left panel) and *CDKN1A* (right panel) relative mRNA levels measured by RT-qPCR in HEK-293T cells treated as in C. (E) Luciferase activity assay in HEK-293T cells transiently transfected with different plasmids: luciferase control plasmid without promoter sequences (Control-L), reporter of the *VCAN* promoter’s activity (VCAN-L), hyperactive form of MEK5 (MEK5DD) plus a WT ERK5 (ERK5) or an inactive ERK5 (ERK5-KD) (left panel). Protein expression was measured by western blot, using HA antibody for ERK5 levels and Vinculin as loading control (right panel). (F) Luciferase activity assay in SK-LMS-1 cells transiently transfected with the same plasmids as in panel E. (G) SK-LMS-1 cells were transduced with lentiviral vector pBabe Control (pBABE Cont.) or expressing the hyperactive form of MEK5 (MEK5DD), and selected cells were analyzed by western blot against the indicated antibodies (left panel). *VCAN* relative mRNA levels were evaluated by RT-qPCR in these cells (right panel). Graphics represent the mean +/-SD of 3 independent experiments. The unpaired Student’s t-test was used to assess statistical significance. *p<0.05; **p<0.01; ***p<0.001.

### ERK5 regulates *VCAN* promoter activity

To further confirm the possible transcriptional regulation of *VCAN* by the ERK5-dependent signaling pathway, we performed transient transfection experiments in the HEK-293T cell line with a human *VCAN* promoter construct (-900 to +100 bp relative to the genés transcription start site) in a *Renilla reniformis* luciferase reporter gene vector. First, we proved the response of endogenous VCAN to chemical inhibition of ERK5 (Fig. 2D), confirming our previous observations. Next, to trigger activation of the ERK5 pathway we co-transfected HEK-293T cells with expression vectors encoding a constitutively active MEK5 (MEK5DD) and a wild type ERK5. In these experimental conditions, ERK5 activation determined a significant increase in VCAN-luciferase reporter activity, that was not observed in the presence of kinase dead inactive ERK5 (ERK5KD) (Fig. 2E). The same results were obtained in the human SK-LMS-1 cells, where both transient transfection (Fig. 2F) and stable expression of MEK5DD similarly upregulated endogenous *VCAN* expression (Fig. 2G).

In sum, all lines of evidence support the ERK5 signaling pathway as a major regulator of *VCAN* expression at the transcriptional level.

### VCAN mediates *in vitro* and *in vivo* biological effects associated with the ERK5 signaling pathway

Next, we evaluated the biological role of the ERK5-VCAN signaling axis in our experimental sarcoma models. Using specific shRNAs targeting *MAPK7* or *VCAN* (Fig. 2A and 3A), we observed similar effects in SK-LMS-1 cells regarding cell growth (Fig. 3B) and colony formation (Fig. 3C). Consistently, in both AA and EC experimental models we obtained nearly identical results (Supplementary Fig. 3 and 4). Another biological effect known to be controlled by ERK5 is cell migration (32). Initially, to avoid the proliferation effects on wound healing assays in our model of SK-LMS-1, we tested low serum conditions with or without the growth inhibitor Mimosine, confirming comparable results in both conditions (Supplementary Fig. 5). Under these experimental setting the knockdown of ERK5 or VCAN resulted in a significant reduction in cell migration ability (Fig. 4A). In addition to migration, adhesion has also been related to ERK5 (33). Consistent with this, lack of ERK5 or VCAN expression rendered a decrease in adhesion to collagen-coated wells (Fig. 4B). We then assessed tumor growth *in vivo* using SK-LMS-1 xenografts which demonstrated that VCAN or ERK5 knockdown similarly increased tumor latency (Fig. 4C). Furthermore, small tumors derived from ERK5 or VCAN knockdown cells showed a clear recovery of *MAPK7* and *VCAN* expression (Supplementary Fig. 6A) with identical histological features (Supplementary Fig. 6B), indicating that the observed small tumors likely arise from poorly interfered (escape) cells, supporting the critical role of ERK5 and VCAN in the *in vivo* tumor growth of our experimental model.

**Figure 3.**
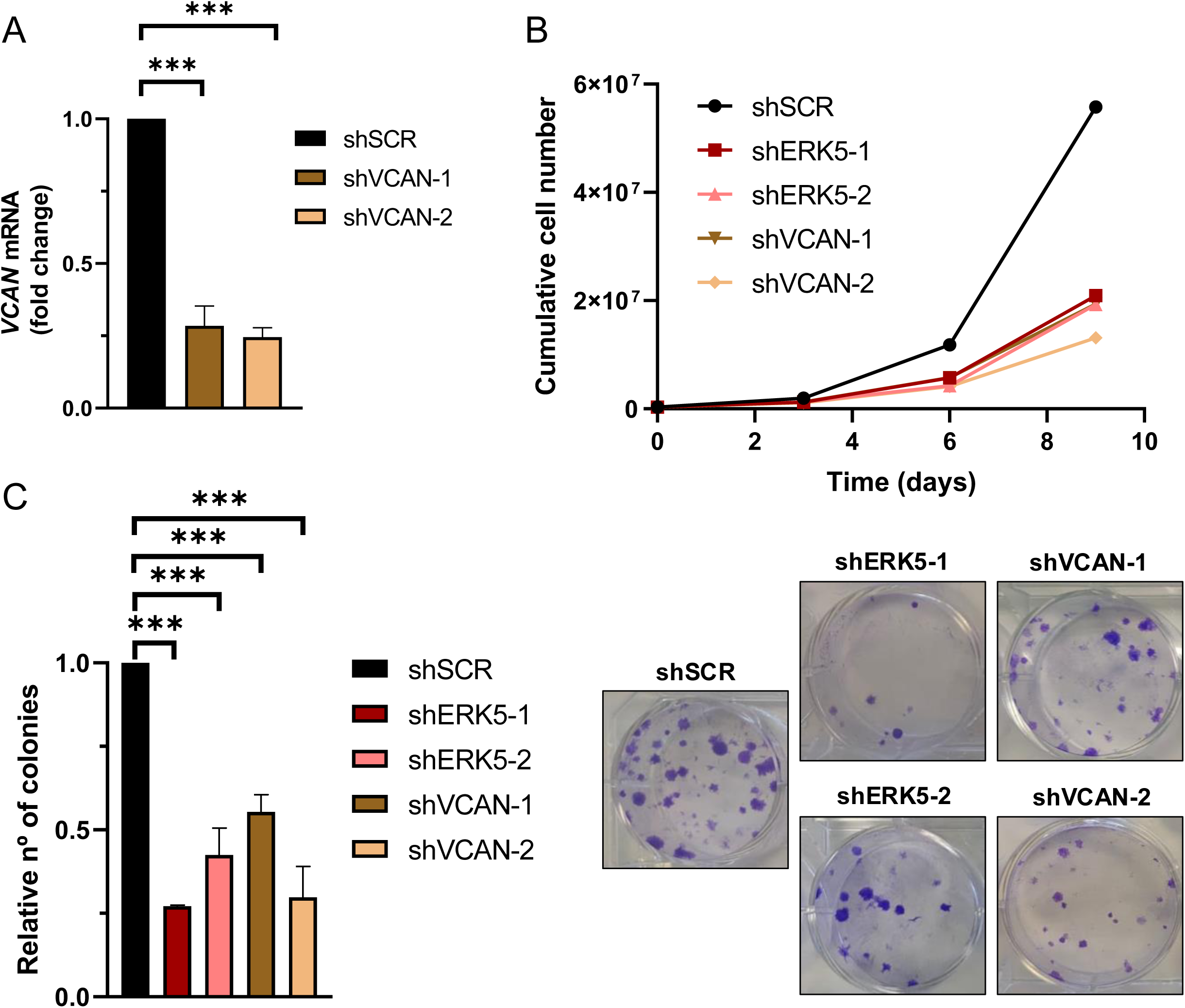
ERK5 and VCAN mediate proliferation and colony formation in SK-LMS-1 cells. (A) *VCAN* relative mRNA levels were evaluated by RT-qPCR in SK-LMS-1 cells infected with lentiviruses carrying PLKO.1-shScramble vector (shSCR) or PLKO.1-shRNAs for VCAN (shVCAN-1/2). (B) Growth curves of 3 × 10^5^ shSCR, shERK5-1/2 or shVCAN-1/2 SK-LMS-1 cells seeded in 100 mm plates and replated every 3 days up to day 9. Representative experiment out of 3 from different pools of infections with nearly identical results. (C) Relative number of colonies calculated by clonogenic assays with 200 cells/well of shSCR, shERK5-1/-2 and shVCAN-1/-2 SK-LMS-1 cells stained with crystal violet after 12 days (left panel). Representative images of clonogenic assays from different conditions (right panel). Histograms represent the mean +/-SD of 3 independent experiments from different pools of infection. The unpaired Student’s t-test was used to assess statistical significance. ***p<0.001.

**Figure 4.**
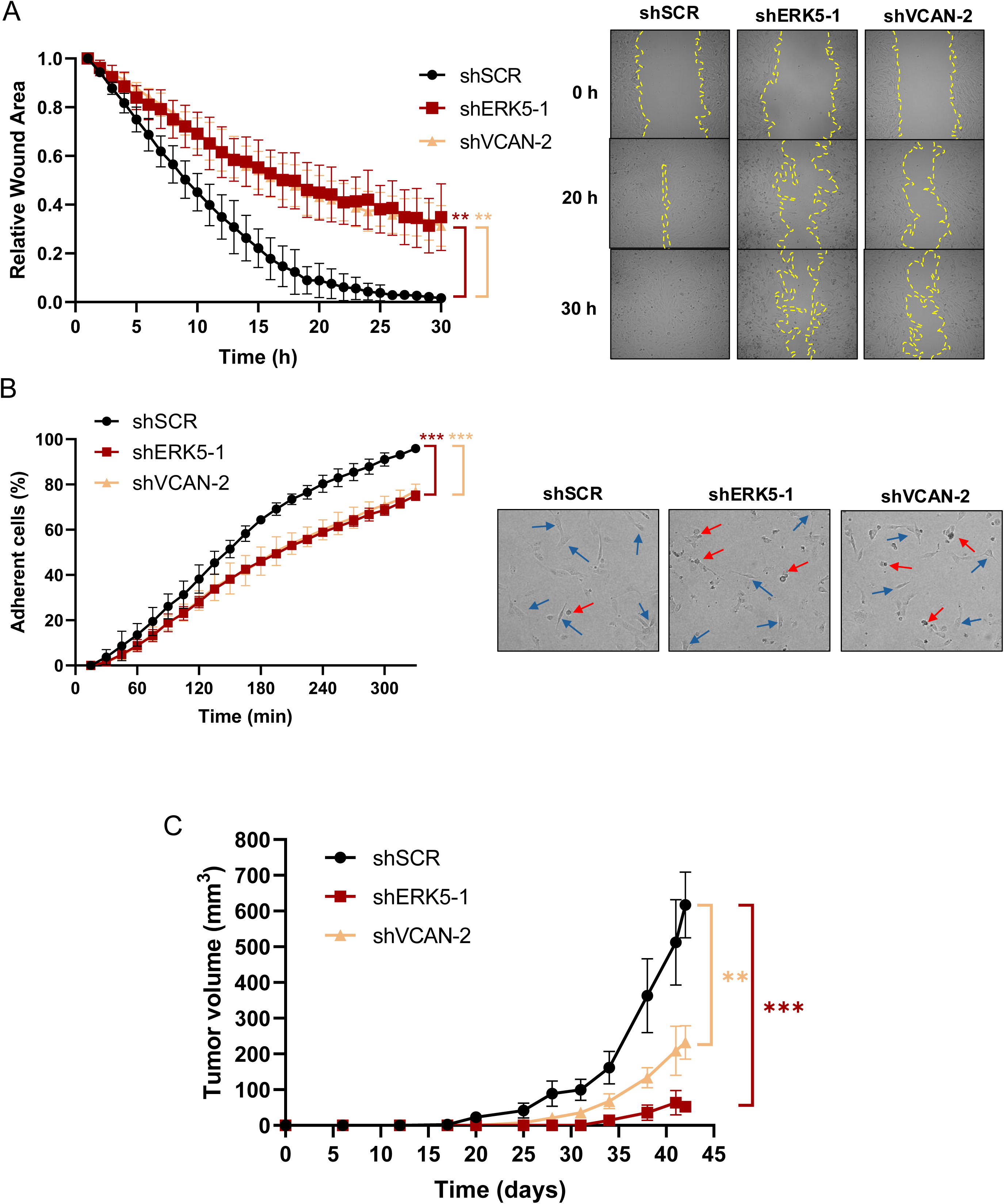
ERK5 and VCAN are involved in *in vitro* migration and adhesion, and *in vivo* tumor growth capabilities of SK-LMS-1 cells. (A) Wound healing assays were performed in SK-LMS-1 cells infected with lentiviruses carrying PLKO.1-shScramble (shSCR), PLKO.1-shRNA ERK5-1 (shERK5-1) or PLKO.1-shRNA VCAN-2 (shVCAN-2) vectors. Left panel shows the mean +/-SD of relative wound area of shSCR, shERK5-1 and shVCAN-2 SK-LM-S1 cells followed up to 30 hours, from 3 independent pools of infection. Right panel shows representative images from one experiment. (B) Percentage of shSCR, shERK5-1 and shVCAN-2 SK-LMS-1 cells fully adhered until 330 minutes after seeding (left panel). Graphic represents the mean +/-SD of 3 independent experiments from different pools of infection. Right panels show representative images of cells taken 300 minutes after seeding. Blue arrows mark fully adherent and expanded cells, red arrows mark not attached cells. (C) tumor growth of 2 × 10^6^ subcutaneously inoculated shSCR, shERK5-1 and shVCAN-2 SK-LM-S1 cells in NSG mice (n=4). The graphic represents the mean ± SEM. The unpaired Student’s t-test was used to assess statistical significance. **p<0.01; ***p<0.001.

All the previous data showed a strong correlation between ERK5- and VCAN-dependent biological outcomes and expression; however, they did not establish a cause-effect relationship. To address this issue, we generated SK-LMS-1 cells overexpressing exogenous *VCAN* (Fig. 5A). In a *VCAN* overexpression context, ERK5 silencing did not modify *VCAN* expression (Fig. 5B), had a discrete effect on proliferation (Fig. 5C) and foci formation (Fig. 5D and Supplementary Fig. 7A), and no effect on migration (Fig. 5E, Supplementary Fig. 7B) and adhesion (Fig. 5F, Supplementary Fig. 7C). Importantly, reduced ERK5 levels did not modify the *in vivo* tumorigenicity of *VCAN* overexpressing SK-LMS-1 cells (Fig. 5G). Furthermore, analysis of endpoint tumors showed a full recovery of *MAPK7* expression and no effect on *VCAN* overexpression as well as an identical histology (Fig. 5H and I), supporting the critical role of VCAN in ERK5-dependent tumorigenicity.

**Figure 5.**
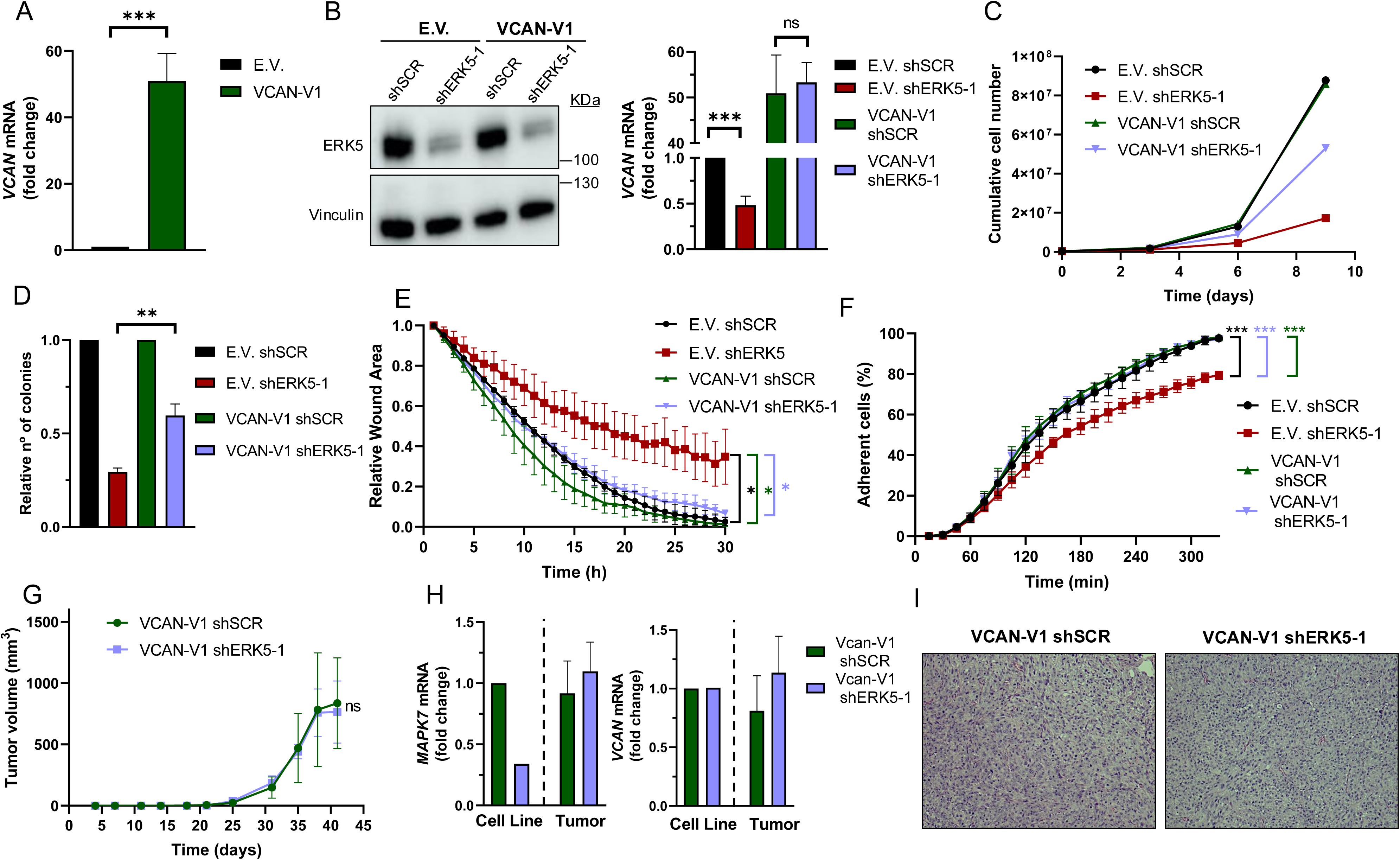
VCAN overexpression rescues the effects of ERK5 silencing in SK-LMS-1 cells. (A) *VCAN* relative mRNA levels of SK-LMS-1 cells transfected with empty vector pSecTag A (E.V.) and pSecTagA VCAN-V1 (VCAN-V1), analyzed by RT-qPCR. (B) ERK5 protein levels of E.V. and VCAN-V1 SK-LMS-1 cells infected with lentiviruses carrying PLKO.1-shScramble (shSCR) or the human PLKO.1-shRNA ERK5-1 (shERK5-1) vectors were evaluated by western blot. Vinculin was used as a loading control (left panel). *VCAN* mRNA levels of the same cells were analyzed by RT-qPCR (right panel). (C) Growth curves of 3 × 10^5^ E.V. shSCR, E.V. shERK5-1, VCAN-V1 shSCR and VCAN-V1 shERK5-1 SK-LMS-1 cells in 100 mm plates. Every 3 days, cells were counted and replated in the same manner up to day 9. The graphic shows the cumulative cell number from a representative experiment out of 3 with nearly identical results in different pools of infections. (D) Relative number of colonies obtained in clonogenic assays of SK-LMS-1-derived cell lines E.V. shSCR, E.V. shERK5-1, VCAN-V1 shSCR and VCAN-V1 shERK5-1. Graphics represent the mean +/-SD of 3 independent experiments from different pools of infection. (E) Relative wound area of SK-LMS-1-derived cells up to 30 hours after wound was made. Graphics represent the mean +/-SD of 3 independent experiments from different pools of infection. (F) Percentage of SK-LMS-1-derived cells fully adhered to the surface up to 330 minutes after seeding. Graphics represent the mean +/-SD of 3 independent experiments from different pools of infection. (G) Tumor growth of 2 × 10^6^ VCAN-V1 shSCR or VCAN-V1 shERK5-1 SK-LMS-1-derived cell lines subcutaneously injected in NSG mice (n=4) at the indicated times. The graphic represents the mean ± SEM for each timepoint. (H) *MAPK7* and *VCAN* relative mRNA levels of VCAN-V1 shSCR or VCAN-V1 shERK5-1 SK-LMS-1-derived cell lines before injection (Cell Line) and from recovered tumors (Tumor) analyzed by RT-qPCR. I) Representative images of hematoxylin and eosin staining of tumors obtained from SK-LMS-1-derived cell lines. Pictures are shown at 20X magnification. The unpaired Student’s t-test was used to assess statistical significance. *p<0.05 ; **p<0.01; ***p<0.001.

In sum, all the above data demonstrate that VCAN regulation is a critical event in the biological processes governed by the ERK5 signaling pathway in sarcoma biology.

### VCAN influences the ERK5-dependent transcriptional landscape

Next, we decided to evaluate the effect of the signaling axis ERK5-VCAN at the transcriptional level. For this purpose, SK-LMS-1 cells were effectively transduced with lentiviral vectors carrying specific shRNAs for *MAPK7* or *VCAN* and analyzed by RNA-seq. ERK5 abrogation modulated 822 genes (423 upregulated and 399 downregulated). As expected, DEGs included *MAPK7* and *VCAN*, while other members of the proteoglycan family were either not expressed or unaffected (Fig. 6A). The DEGs were associated with established biological functions of ERK5 (Fig. 6B). On the other hand, *VCAN* knockdown resulted in 1,321 DEGs (512 upregulated and 809 downregulated, Fig. 6C), showing a pattern consistent with the expected biological role of VCAN, such as response to growth factors, wound healing or adhesion (Fig. 6D). A comparison of DEGs between ERK5 and VCAN knockdowns revealed a significant overlap (Fisher’s exact test, OR = 3.89, p-value = 3.92 × 10⁻⁴⁵), indicating that the 200 shared genes greatly exceed the number expected by chance (Fig. 6E). Moreover, the magnitude of gene regulation log fold change (LogFC) in both conditions showed a moderate but significant correlation for both upregulated (R = 0.53, p < 0.001) and downregulated genes (R = 0.42, p < 0.001) (Supplementary Fig. 8). Functional enrichment analysis of these DEGs revealed that 91 GO categories were common to both ERK5 and VCAN interference (Fig. 6F). Interestingly, when focusing on the enriched functions within the 200 overlapping

**Figure 6.**
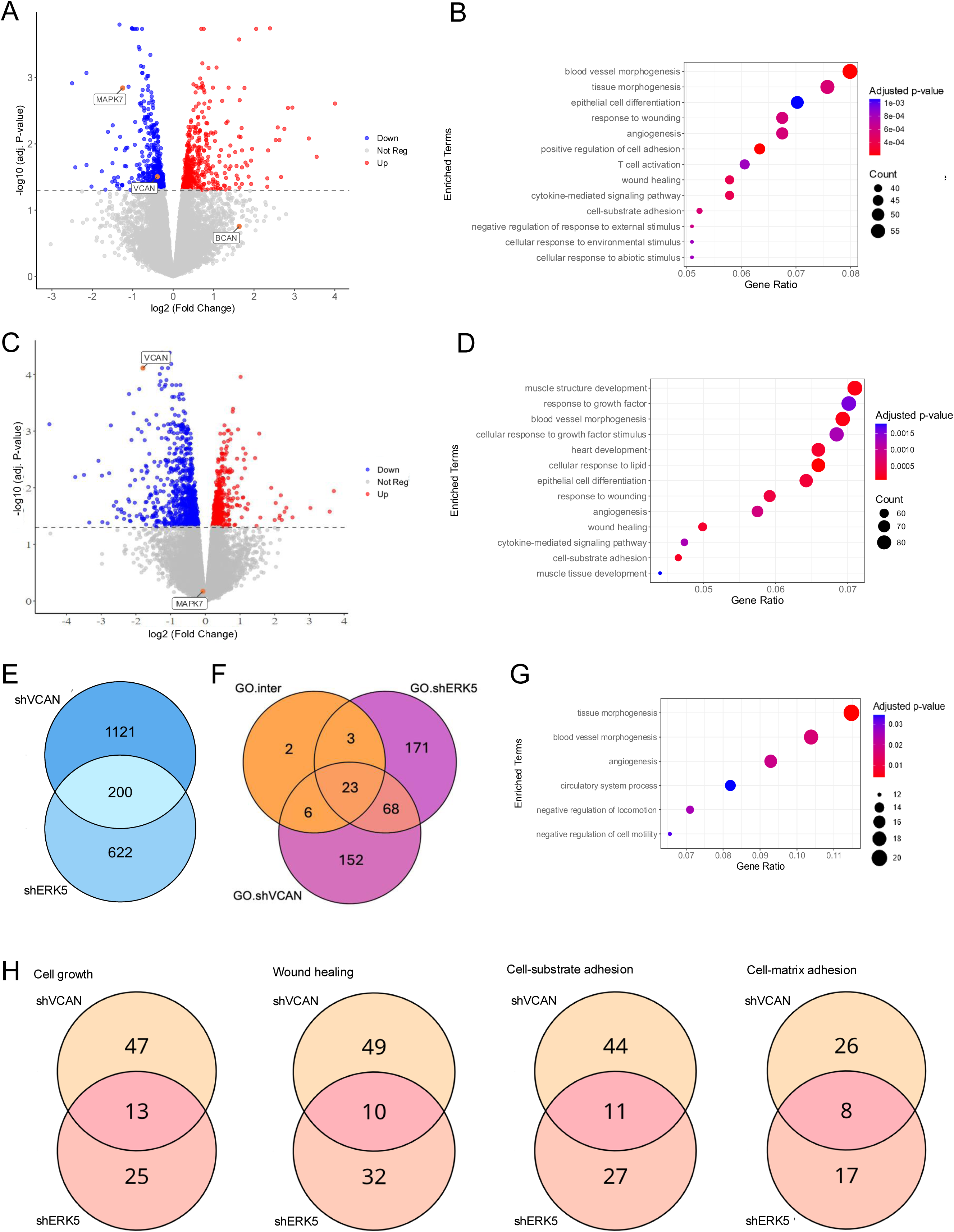
Transcriptional analysis of the ERK5-VCAN signaling axis. (A) Volcano plot showing differential gene expression after *MAPK7* suppression. Genes with significant upregulation (adj. p-value < 0.05 and log2(Fold Change) > 0) are highlighted in red, while significantly downregulated genes (adj. p-value < 0.05 and log2(Fold Change) < 0) are shown in blue. (B) Gene Ontology Biological Process (GO-BP) enrichment analysis of significantly regulated genes after *MAPK7* suppression, showing the top 13 enriched terms with the highest Gene Ratio. (C) Volcano plot of gene expression changes after VCAN suppression, using the same statistical criteria as in panel A. (D) GO-BP enrichment analysis for genes significantly regulated by *VCAN* suppression, highlighting the top 13 enriched terms by Gene Ratio. (E) Venn diagram showing the overlap of genes regulated by both *MAPK7* and *VCAN* suppression. (F) Venn diagram showing enriched GO-BP specific to *MAPK7* suppression (GO.shERK5), *VCAN* suppression (GO.shVCAN), and those common to both (GO.inter). (G) GO-BP enrichment analysis of genes regulated by *VCAN* and *MAPK7* knockdown focusing on terms related to vasculature development and cell motility. (H) Venn diagrams showing the overlap of genes within selected GO-BP enriched by both *MAPK7* and *VCAN* abrogation.

DEGs, only 23 terms were shared (Fig. 6F), most notably those related to vasculature development and cell motility (Fig. 6G). Further analysis of the 91 GO categories common to both ERK5 and VCAN knockdowns, including functions such as cell growth, wound healing, cell adhesion to substrates or to the extracellular matrix, revealed that VCAN contributes to approximately 30% of the transcriptional regulation mediated by ERK5 (Fig. 6H and Supplementary Fig 9). This also suggests, however, that ERK5 suppression can impact those functional categories through mechanisms independent of VCAN regulation.

### VCAN expression in human sarcoma samples correlates with ERK5 expression

Finally, we sought to extrapolate our findings to a clinical context. To begin with, we performed *in silico* analysis of the expression levels of the chondroitin-sulfate proteoglycan family members (VCAN, ACAN, BCAN and NCAN) using published datasets (34). As it is shown in Supplementary Fig. 10, data from the TGCA series for STS revealed that VCAN displayed significantly a higher expression compared to other family members, highlighting the VCAN’s unique prominence in STS biology across chondroitin-sulfate proteoglycan family members.

Furthermore, the TCGA-SARC cohort displayed some of the highest average expression levels of *MAPK7* and *VCAN* genes across the entire dataset, suggesting a potential association between the expression of these two genes (Supplementary Fig. 11 A and B). To further investigate this relationship, we analyzed an independent cohort of 216 patients with available matched normal and tumor tissue after quality control process out of a total of 222 patients (Supplementary Table 3). All patients underwent surgical resection at a single institution. RNA sequencing was performed on tumor and paired adjacent normal tissues. In this independent cohort, we observed a pronounced upregulation of both *MAPK7* and *VCAN* in tumor samples compared to matched normal tissue across multiple STS histologies arising in the extremities (Fig. 7). A similar trend was also observed in retroperitoneal STS samples, although to a lesser extent likely due to the expected smaller number of available cases for certain histologies, (Supplementary Fig. 11 C and D). Of note, LMS and UPS, which had a marked upregulation in *MAPK7* and *VCAN*, are the two STS histologies previously shown to be significantly dependent on ERK5 signaling in preclinical murine sarcoma models (8,9). Based on this, we conducted independent analyses of these subtypes and found a robust and statistically significant correlation between *MAPK7* and *VCAN* mRNA expression levels in both LMS (n=21, r=0.58) and UPS (n=25, r=0.85), irrespective of tumor localization (Fig. 8A and B). In addition, immunohistochemical studies of a small independent cohort of LMS (n=10) and UPS (n=9) (see Supplementary Table 4) from a different institution revealed a marked correlation between ERK5 and VCAN protein expression (Fig 8C and D).

**Figure 7.**
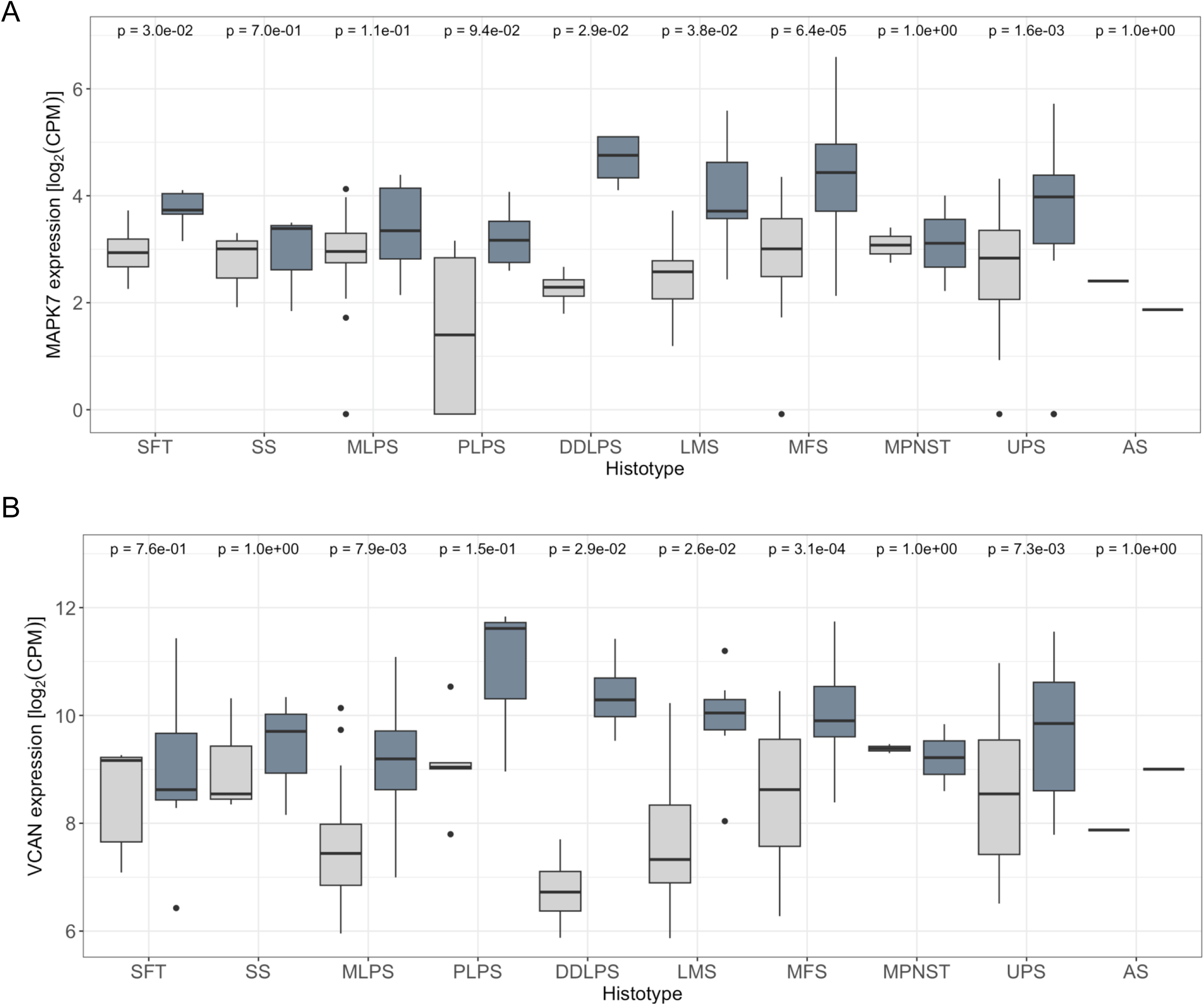
*MAPK7* and *VCAN* mRNAs are overexpressed in soft tissue sarcomas. A) Comparison of *MAPK7* mRNA expression levels (log₂ CPM) between extremity softtissue sarcomas and paired normal tissue, using RNA-Seq data from the SARCOMICS study. B) Comparison of *VCAN* mRNA expression levels (log2 CPM) in the same collection of extremity soft-tissue sarcomas and paired normal tissue by RNA-Seq. Wilcoxon test was used to address statistical significance. WDLPS: Well-differentiated liposarcoma; DDLPS: dedifferentiated liposarcoma; PLPS: pleomorphic liposarcoma; MLPS: myxoid liposarcoma; LMS: leiomyosarcoma; MPNST: malignant peripheral nerve sheath tumor; UPS: undifferentiated pleomorphic sarcoma; SS: synovial sarcoma; AS: angiosarcoma; SFT: solitary fibrous tumor; MFS: myxofibrosarcoma.

**Figure 8.**
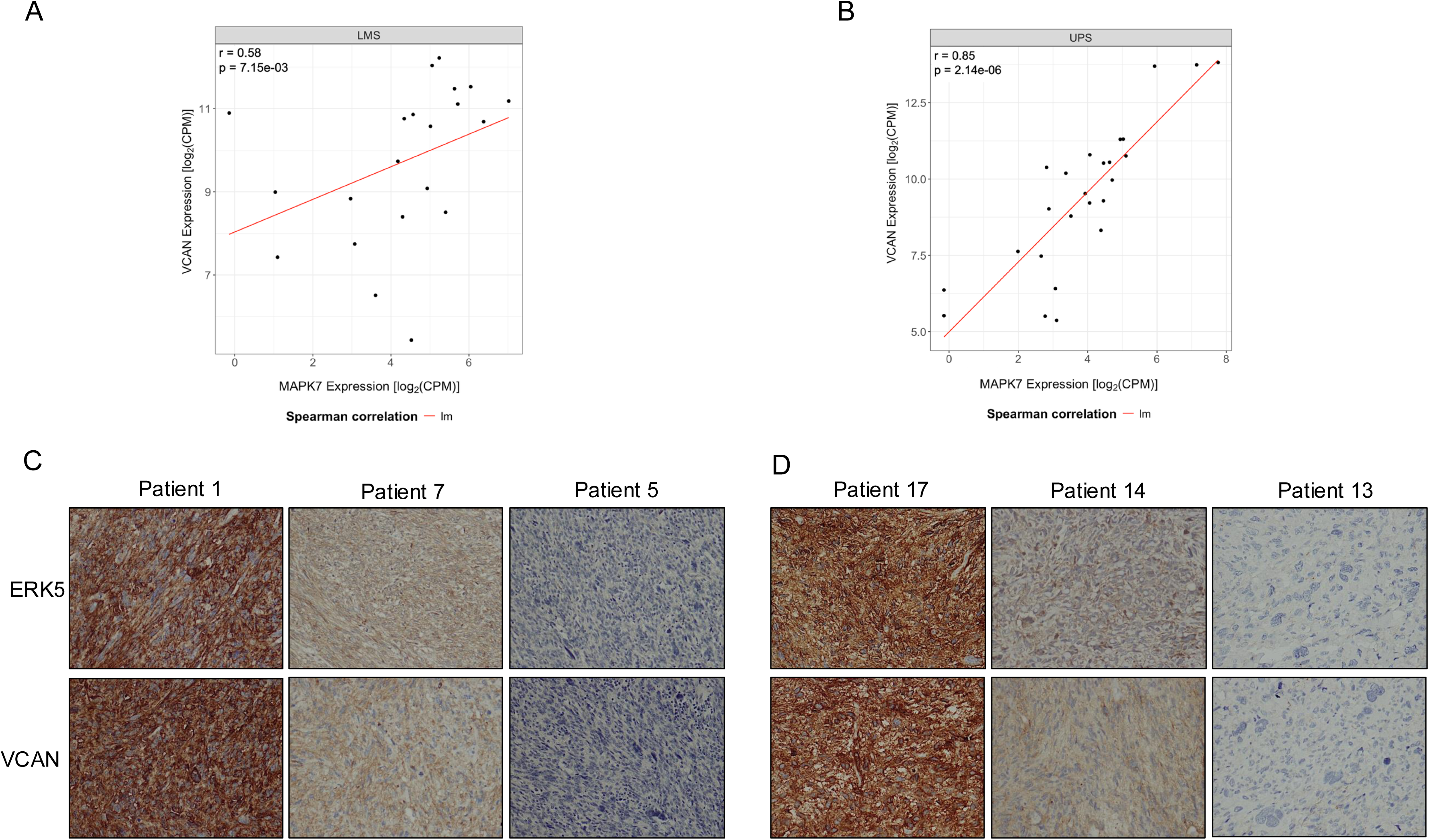
*MAPK7* and *VCAN* gene expression correlate in leiomyosarcoma (LMS) and undifferentiated pleomorphic sarcoma (UPS). (A) and (B) Scatter plots showing the correlation between *MAPK7* and *VCAN* gene expression in LMS and UPS samples combining together extremity and retroperitoneal sarcoma from the SARCOMICS study, with linear regression trend line depicted in red, Spearman’s correlation coefficient (r) and corresponding p-values indicated within each panel. The analysis was performed for 21 LMS (A) and 25 UPS (B). (C) Representative images of immunohistochemical staining for ERK5 and VCAN in 10 LMS samples and 9 UPS (D) samples from another independent cohort.

In summary, this evidence achieved on samples from patients with STS is consistent with our *in vitro* experiments, supporting that the ERK5 signaling pathway regulates VCAN expression in human tumors.

## 4. Discussion

Increased expression/activation of components of the ERK5 route has been linked to the initiation and progression of several types of tumors (35). However, the mechanisms by which this MAPK pathway contributes to the oncogenic phenotype are still unclear. In this paper, we identify VCAN as a critical mediator in some of these prooncogenic actions, including cell proliferation, migration, adhesion and, more importantly, *in vivo* tumorigenesis. The preclinical evidence, together with the correlation between ERK5 and VCAN in patient samples, opens new possibilities to be considered for the therapy of tumors in which the ERK5-VCAN axis may play a role in their pathophysiological progression.

Several solid data support a link between ERK5 and VCAN. First, the discovery of *VCAN* as a new transcriptional target of the ERK5 signaling pathway. Therefore, *VCAN* could be included in the list of genes to be used as biomarkers of the ERK5 route, similar to other previously identified targets such as the cell cycle regulators p21 or p27 (31,36) or, more recently, metabolic enzymes as PFKFB3 and glutaminase (37,38). Furthermore, our observation seems to have a wider character not restricted to mesenchymal tumors such as STS, since we have observed such a relationship in cancer derived cell lines of epithelial origin as renal or breast cancer. This may have relevant implications for the biomarking of tumors in which the ERK5 pathway is upregulated and therefore susceptible to be manipulated with therapeutic purposes. The known mechanisms of *VCAN* regulation are complex and depend on the cellular context. Since the initial isolation of the VCAN proximal promoter, putative binding sites for the transcription factors SP1, AP2, C/EBP and CTF/CBF were identified (39). Subsequently, it was shown that in human melanoma-derived cell lines the binding of the transcription factors SP1 and TCF-4 is responsible for most of the activity of the *VCAN* proximal promoter (40). The TCF-4/β-catenin signaling pathway is also key for the activation of the *VCAN* promoter in vascular smooth muscle cells (41) and in dermal papilla cells (42). Of note, other important stimuli and proteins regulate *VCAN* expression, such as hypoxia (43), androgen receptor (44), activin A (45), or FoxQ1 (46), among many others. For example, recent evidence has linked VCAN and ERK1/2 in colorectal cancer; however, ERK5 was not analyzed in this report (47). Therefore, the identification of the transcription factors responsible for the crosstalk between ERK5 and VCAN will require future detailed investigations and is likely to depend on distinct factors depending on the cellular context.

From a biological perspective, the identification of *VCAN* expression as a target of ERK5 signaling has several important implications. As mentioned above, the ERK5 pathway has been proposed as a key mediator of several aspects of tumor progression, with direct involvement in processes such as migration, invasion, angiogenesis, etc. (for a review see (35)) that have also been associated with VCAN (48). For example, recent work links ERK5 to cell adhesion via FAK (33), which is remarkable given that FAK has also been identified as a target of VCAN (49,50). Similarly, migration has been extensively studied in the context of ERK5 (33,51) and VCAN (52,53), reinforcing the connection between both molecules. All these observations align well with the context of epithelial-mesenchymal transition, in which ERK5 and VCAN play well-defined roles, often acting in concert with factors such as TGF-β1 and Snail (54–56).

Regarding LMS our data support previous findings on the role of VCAN in this particular type of STS (57). In addition, this signaling axis could be considered to explain characteristics as the high metastatic potential of retroperitoneal LMS, with a vascular origin and a high risk of distant metastasis in approximately 50% of the cases (58,59). This observation could be extended to UPS, a histology with a high risk of developing distant metastases after surgery (60).

An important implication of our findings is their potential impact on cancer therapy. In this regard, VCAN has been related to the tumor response to conventional chemo/radiotherapy (61–63) in which ERK5 has also been implicated (64–66). Additionally, the axis ERK5-VCAN may influence other therapeutic strategies. For example, it has been recently reported that ERK5 signaling mediates cellular responses to death-receptor agonists (67) in which VCAN also plays a role (68,69). Furthermore, immunotherapy, which has become one of the most promising tools for controlling tumor growth and progression, could similarly be affected by this signaling axis. Recent evidence demonstrates that ERK5 inhibition enhances the efficacy of PD-1-based therapy by modulating TGFβ1 signaling (70), and also by regulating key immune response molecules (71,72). Similarly, VCAN has emerged as a critical determinant in immunotherapy response (62,73), for example, by controlling T-cell trafficking (74). In this context, regulating VCAN expression via ERK5 could play a key role in advancing immunotherapeutic strategies for sarcomas such as UPS, which has shown promising responses to immunotherapy in clinical trials at both early (SARC032 (75)) and metastatic (SARC028 (76)) stages. In fact, VCAN has been proposed as a novel immunotherapeutic target (77), further underscoring its therapeutic relevance that could avoid undesirable effects associated with therapy based on ERK5 inhibition (78).

In summary, our results establish VCAN as a new downstream effector of the ERK5 signaling pathway, playing a critical role in mediating its biological functions in STS pathophysiology and paving the way to new therapeutic opportunities. Further investigation is required to determine whether these observations are applicable to other tumor types and to identify the interacting proteins that regulate the ERK5-VCAN signaling axis.

## Acknowledgements

This work has been supported by grant PID2021-122222OB-I00 and PID2019-104416RB-I00 funded by MCIN/AEI /10.13039/501100011033/ and by FEDER A way to make Europe to RSP and JCR-M. RSP and MJRH are also funded by UCLM with grant 2022-GRIN-34150 and by Agencia de Investigación e Innovación, Junta de Comunidades de Castilla-La Mancha grant SBPLY/23/180225/000007. SP is supported by “5×1000 Founds” – 2016, Italian Ministry of Health – Institutional Grant BRI2017 from Fondazione IRCCS Istituto Nazionale dei Tumori di Milano. SP and CS are supported by AIRC Individual Grant - Next Gen Clinician Scientist “Fondazione 13 marzo” [ID#28546]. YBD is supported by Grant PID2020-118821RB-I00 funded by MCIN/AEI/10.13039/501100011033 and by FSE. FJC is funded by grant SBPLY/23/180225/000007. JJS is funded by “Contrato predoctoral para la formación de personal investigador en el marco del Plan Propio de I+D+I UCLM 2020-PREDUCLM-15144, co-financed by the European Social Fund Plus (ESF+). ADC is funded with FPU fellowships from the Spanish Ministry of Education. Work in AP and AE-O laboratories were funded by Ministry of Economy and Competitiveness of Spain (PID2020-115605RB-I00), the Instituto de Salud Carlos III through CIBERONC, and grant number PI19/00840 cofunded by the European Union, the CRIS Cancer Foundation and the Regional Development Funding Program (FEDER) “A way to make Europe”. Work in ER laboratory was funded by Associazione Italiana per la Ricerca sul Cancro (AIRC) and Fondazione CR Firenze (IG 2018 - ID. 21349 project). We are very grateful for the funds provided by Fundación Leticia Castillejo, Taller Solidario Árbol De La Vida (Las Pedroñeras), Asociación Comarcal Contra El Cáncer De Motilla Del Palancar, Asociación Jareña Contra el Cáncer. We also would like to thank the staff from the Biobank at the Complejo Hospitalario Universitario de Albacete for their technical Support.

## Data Availability Statement

All data and materials are available upon reasonable request. RNA-seq analysis data for SK-LMS-1 cells with abrogated ERK5 or VCAN expression have been deposited in the Gene Expression Omnibus (GEO) under the accession code GSE289613.

## Authorship contribution statement

Jaime Jiménez-Suárez: Methodology, Investigation, Visualization, Writing-Reviewing Editing.

Francisco J. Cimas: Methodology, Visualization, Investigation, Supervision, Writing-Reviewing Editing, Validation.

José Joaquín Paricio: Methodology, Investigation, Visualization, Writing-Reviewing Editing.

Borja Belandia: Methodology, Investigation, Supervision, Visualization, Writing-Reviewing Editing.

Yosra Berrouayel: Data Curation, Formal Analysis, Visualization, Writing-Review Editing.

Elena Arconada-Luque: Methodology, Investigation, Visualization.

Sofía Matilla-Almazán: Methodology, Investigation, Visualization.

Cesare Soffientini: Formal Analysis, Investigation, Data Curation.

Stefano Percio: Formal Analysis, Investigation, Data Curation.

Silvia Redondo-García: Methodology, Investigation.

Natalia Garcia-Flores: Methodology, Investigation.

Cristina Garnés-García: Methodology, Investigation.

Pablo Fernández-Aroca: Methodology, Investigation.

Juan Jesus Martínez-Gomez: Methodology, Investigation.

Juan Paricio-Canada: Methodology, Investigation.

Syong Hyun Nam Cha: Methodology, Visualization, Writing-Reviewing, Editing.

Antonio Fernández Aramburo: Conceptualization, Writing-Reviewing, Editing.

Elisabetta Rovida: Conceptualization, Writing-Reviewing, Editing.

Juan Carlos Rodríguez-Manzaneque: Supervision, Conceptualization, Writing-Reviewing, Editing.

Atanasio Pandiella: Conceptualization, Writing-Reviewing, Editing.

Azucena Esparís-Ogando: Supervision, Conceptualization, Writing-Reviewing, Editing.

Sandro Pasquali: Formal Analysis, Data Curation, Writing-Reviewing, Editing.

Luis del Peso: Supervision, Data Curation, Formal Analysis, Visualization, Writing-Reviewing, Editing.

María José Ruiz-Hidalgo: Conceptualization, Supervision, Visualization, Writing-Reviewing, Editing, Funding Acquisition.

Ricardo Sánchez-Prieto: Conceptualization, Supervision, writing original draft and Reviewing, Editing, Funding acquisition, Project Administration.

## Abbreviations

3MC: 3Methyl-cholantrene
ECM: Extracellular matrix
DEG: Differentially expressed genes
EMT: Epithelial-mesenchymal transition
GO: Gene Ontology
LMS: Leiomyosarcoma
MAPK: Mitogen-activated protein kinase
STS: Soft tissue sarcoma
UPS: Undifferentiated pleomorphic sarcoma.

## Supplementary files

Supplementary Figure 1. ERK5 regulates *VCAN* mRNA levels in AA and EC cell lines. AA (A) and EC (D) cells were infected with lentiviruses carrying PLKO.1-shScramble vector (shSCR) or the human PLKO.1-shRNA ERK5 1/2 vectors (shERK5-1/2). *MAPK7* mRNA levels were evaluated by RT-qPCR (left panel) and protein levels were evaluated by western blot (right panel). Vinculin was used as a loading control. *VCAN* mRNA levels analyzed by RT-qPCR in AA (B) and EC (E) cells infected with shSCR and shERK5-1/2 vectors. AA (C) and EC (F) cells were treated for 18 hours with the ERK5 chemical inhibitors XMD8-92 (5 µM) and JWG-071 (5 µM), and RT-qPCR was performed to analyze *VCAN* (left panels) and *CDKN1A* (right panels) mRNA levels. Graphics represent the mean +/-SD of 3 independent experiments from different pools of infection. The unpaired Student’s t-test was used to assess statistical significance. *p<0.05; **p<0.01; ***p<0.001.

Supplementary Figure 2. ERK5 regulates *VCAN* mRNA levels in 786-O and Hs 578T cell lines. 786-O (A) and Hs 578T (D) cells were infected with lentiviruses carrying PLKO.1-shScramble vector (shSCR) or the human PLKO.1-shRNA ERK5 vector 1/2 (shERK5-1/2). *MAPK7* mRNA levels were evaluated by RT-qPCR. *VCAN* mRNA levels analyzed by RT-qPCR in 786-O (B) and Hs 578T (E) cells infected with shSCR and shERK5-1/2. 786-O (C) and Hs 578T (F) cells were treated for 18 hours with the ERK5 chemical inhibitor JWG-071 (5 µM), and RT-qPCR was performed to analyze *VCAN* (left panels) and *CDKN1A* (right panels) mRNA levels. Graphics represent the mean +/-SD of 3 independent experiments from different pools of infections. The unpaired Student’s t-test was used to assess statistical significance ***p<0.001.

Supplementary Figure 3. ERK5 and VCAN contribute to the regulation of proliferation and colony formation capacity in AA cells. (A) AA cells were infected with lentiviruses carrying PLKO.1-shScramble vector (shSCR) or the human PLKO.1-shRNA VCAN 1/2 vectors (shVCAN-1/2). *VCAN* mRNA levels were evaluated by RT-qPCR. (B) For growth curves, 2.5 × 10^5^ shSCR, shERK5-1/2 or shVCAN-1/2 AA cells were seeded in 100 mm plates. Every 3 days, cells were counted and replated in the same manner up to day 9. The graphic shows the cumulative cell number from a representative experiment out of 3 from different pools of infection with nearly identical results. (C) For evaluation of colony formation capacity, clonogenic assays were performed by seeding 400 cells/well in a 6_ well plate and they were revealed after 12-14 days. Relative number of colonies obtained in clonogenic assays of shSCR, shERK5-1/2 and shVCAN-1/2 AA cells (left panel). Representative images of clonogenic assays from the different cell lines (right panel). The graphic represents the mean +/-SD of 3 independent experiments from different pools of infections. The unpaired Student’s t-test was used to assess statistical significance. **p<0.01; ***p<0.001.

Supplementary Figure 4. ERK5 and VCAN contribute to the regulation of proliferation and colony formation capacity in EC cells. (A) EC cells were infected with lentiviruses carrying PLKO.1-shScramble (shSCR) or the human PLKO.1-shRNA VCAN-2 (shVCAN-2) vectors. *VCAN* mRNA levels were evaluated by RT-qPCR. (B) For growth curves, 7.5 × 105 shSCR, shERK5-1 or shVCAN-2 EC cells were seeded in 100 mm plates. Every 3 days, cells were counted and replated in the same manner up to day 9. Graphic shows the cumulative cell number from a representative experiment out of 3 from different pools of infection with nearly identical results. (C) For evaluation of colony formation capacity, clonogenic assays were performed by seeding 600 cells/well in a 6-well plate 6, and they were revealed after 12-14 days. Relative number of colonies obtained in clonogenic assays of shSCR, shERK5-1 and shVCAN-2 EC cells (left panel). Representative images of clonogenic assays from the three different cell lines (right panel). The graphic represents the mean +/-SD of 3 independent experiments from different pools of infections. The unpaired Student’s t-test was used to assess statistical significance. ***p<0.001.

Supplementary Figure 5. Setting the conditions of wound healing assay in SK-LMS-1 cells. Relative wound area of SK-LMS-1 cells with 0,5% FBS concentration with/without mimosine treatment (400 µM) (left panel). Representative images from one out of 3 independent experiments (right panel).

Supplementary Figure 6. Analysis of the tumors obtained in figure 4C. (A) *MAPK7* (left panel) and *VCAN* (right panel) mRNA levels of SK-LMS-1 cells before injection (Cell Line) and from recovered tumors (Tumor), analyzed by RT-qPCR. The unpaired Student’s t-test was used to assess statistical significance. (B) Representative images of hematoxylin and eosin histological study of tumors obtained from SK-LMS-1 derived cell lines, showing lesions with high cell density, spindle-shaped cells or occasionally with epithelioid appearance arranged in disorganized bundles. tumor cells also had vesicular nuclei with macronucleoli, extensive eosinophilic cytoplasm and atypical mitoses in all neoplasms. Pictures are shown at a 20X magnification.

Supplementary Figure 7. Images of results presented in Figure 5. (A) Representative images of clonogenic assays from SK-LMS-1 derived cells. (B) Representative images from wound healing assays in SK-LMS-1 derived cells. (C) Representative images of SK-LMS-1 derived cells taken 300 minutes after seeding on a collagen-pretreated surface; blue arrows mark fully adherent and expanded cells; red arrows mark not attached cells.

Supplementary Figure 8. Scatter plot of logFC in gene expression following VCAN versus ERK5 suppression. Each point represents a gene that was differentially expressed in at least one of the two conditions. Genes upregulated in both conditions are shown in red, those downregulated in both are shown in blue. The Pearson correlation coefficients for commonly upregulated and downregulated genes are 0.53 and 0.42, respectively (p < 0.001 for both).

Supplementary Figure 9. Overlap of enriched biological processes between *MAPK7* and *VCAN* interferences. Histogram shows the proportion of overlapping genes in the enriched biological processes shared between *MAPK7* and *VCAN* interferences. For each category, the proportion is calculated by dividing the number of genes identified in the *MAPK7* and *VCAN* deletion enrichments by the total number of genes enriched after *MAPK7* deletion, with *MAPK7* deletion set as the reference: P=G_(MAPK7∩VCAN)/G_MAPK7, where G_(MAPK7∩VCAN) is the number of overlapping genes between *MAPK7* and *VCAN* deletion enrichments and G_MAPK7 is the total number of genes enriched after *MAPK7* deletion. The dashed red line indicates the median overlap ratio across all categories.

Supplementary Figure 10. mRNA expression (log2 TPM) analysis of *VCAN*, *ACAN, BCAN* and *NCAN* expression in samples from TCGA series using TIMER 2.0 platform. Sarcoma series (SARC Tumor) is marked with a black rectangle and the average mRNA level of each gene in sarcoma is marked with a discontinuous black line. Tumor samples depicted in red and healthy samples depicted in blue. Statistical significance of differential expression evaluated using the Wilcoxon test.

Supplementary Figure 11. mRNA expression (log2 TPM) analysis of *MAPK7* (A) and *VCAN* (B) in the TCGA series using TIMER 2.0 platform. Soft Tissue Sarcoma series (SARC Tumor) is marked with a black rectangle and the average *MAPK7* and *VCAN* mRNA levels in sarcoma are marked with discontinuous black lines. Comparison of *MAPK7* (C) and *VCAN* (D) mRNA expression levels (log2 CPM) in retroperitoneal soft-tissue sarcomas and paired normal tissue from the SARCOMICS study by RNA-Seq. Wilcoxon test was used to address statistical significance. WDLPS: Well-differentiated liposarcoma; DDLPS: dedifferentiated liposarcoma; LMS: leiomyosarcoma; MPNST: malignant peripheral nerve sheath tumor; UPS: undifferentiated pleomorphic sarcoma; SFT: solitary fibrous tumor.

