## Supplementary material for "VCAN is essential for ERK5-driven tumorigenesis in soft tissue sarcoma": Supplemnetary files and tables

Supp. Fig. 1

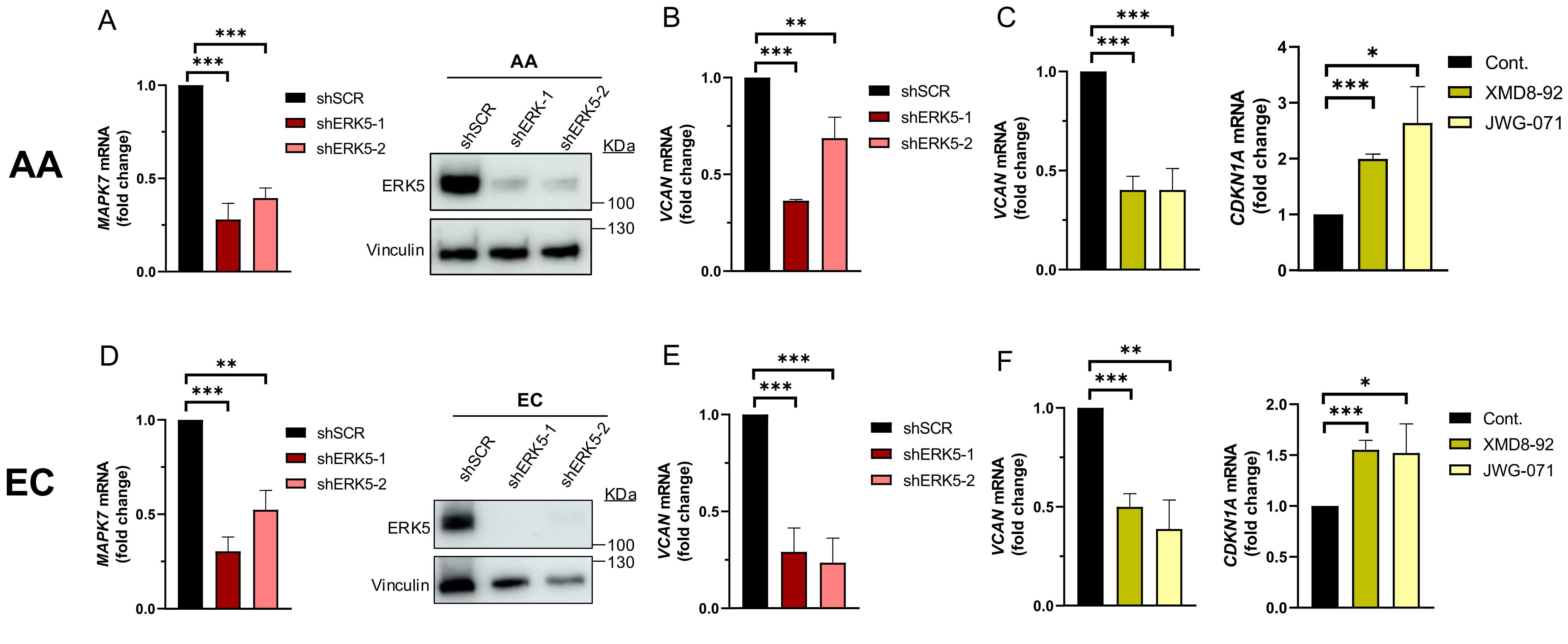
AA
A
***
***
MAPK7 mRNA (fold change)
1.0
0.5
0.0
shSCR
shERK5-1
shERK5-2
AA
shSCR
shERK-1
shERK5-2
KDa
ERK5
100
130
Vinculin
B
**
***
VCAN mRNA (fold change)
1.0
0.5
0.0
shSCR
shERK5-1
shERK5-2
C
***
***
VCAN mRNA (fold change)
1.0
0.5
0.0
*
***
CDKN1A mRNA (fold change)
4
3
2
1
0
Cont.
XMD8-92
JWG-071
EC
D
**
***
MAPK7 mRNA (fold change)
1.0
0.5
0.0
shSCR
shERK5-1
shERK5-2
EC
shSCR
shERK5-1
shERK5-2
KDa
ERK5
100
130
Vinculin
E
***
***
VCAN mRNA (fold change)
1.0
0.5
0.0
shSCR
shERK5-1
shERK5-2
F
**
***
VCAN mRNA (fold change)
1.0
0.5
0.0
*
***
CDKN1A mRNA (fold change)
2.0
1.5
1.0
0.5
0.0
Cont.
XMD8-92
JWG-071

Supp. Fig. 2

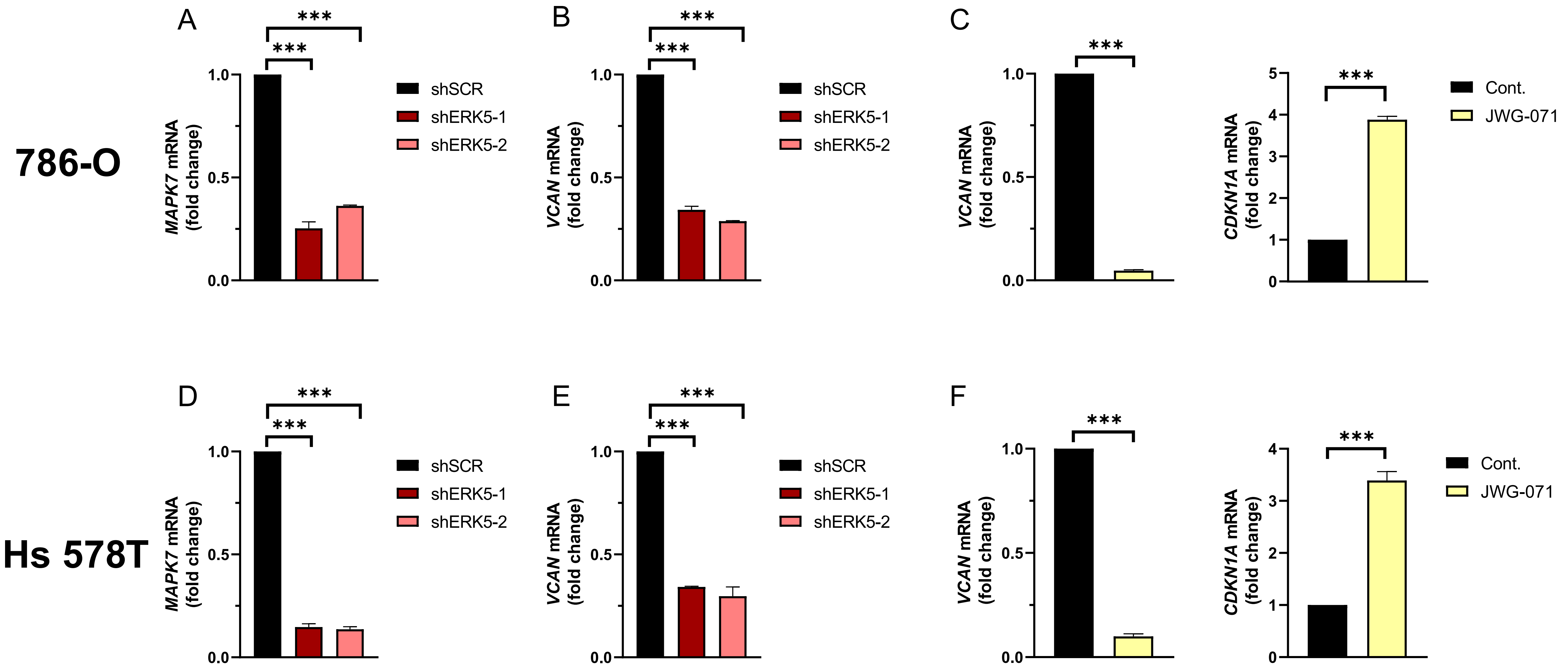

786-O
Hs 578T
A
B
C
D
E
F
***
MAPK7 mRNA (fold change)
VCAN mRNA (fold change)
CDKN1A mRNA (fold change)
1.0
0.5
0.0
5
4
3
2
1
0
shSCR
shERK5-1
shERK5-2
Cont.
JWG-071

### Supp. Fig. 3

## AA

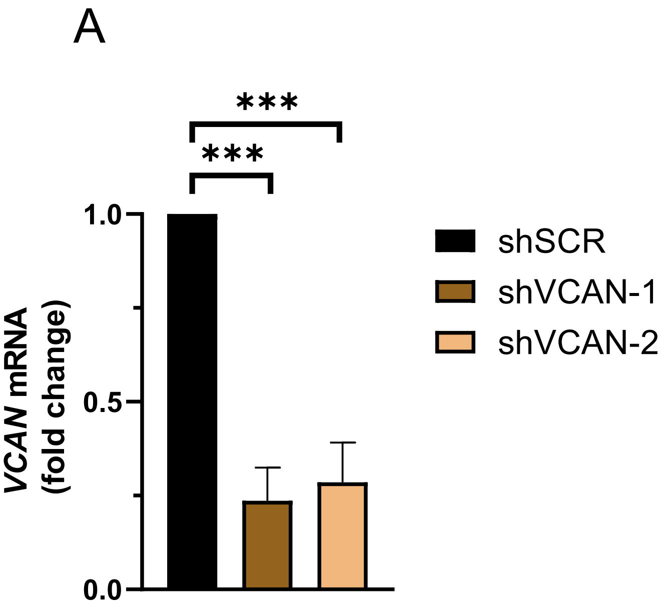

A
***
***
1.0
0.5
0.0
VCAN mRNA
(fold change)
shSCR
shVCAN-1
shVCAN-2


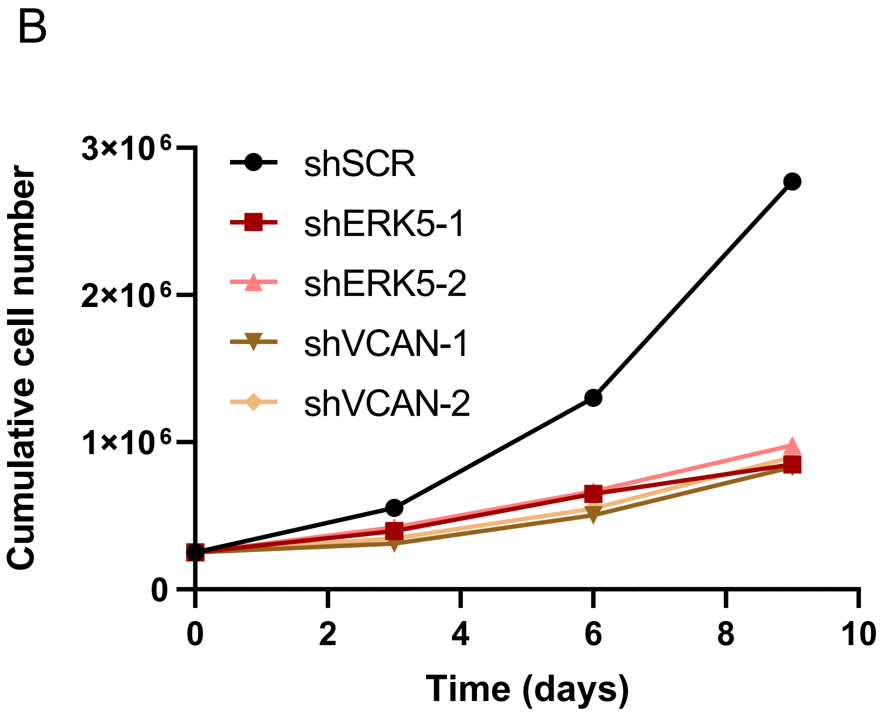

B
3×10⁶
2×10⁶
1×10⁶
0
Cumulative cell number
shSCR
shERK5-1
shERK5-2
shVCAN-1
shVCAN-2
0
2
4
6
8
10
Time (days)


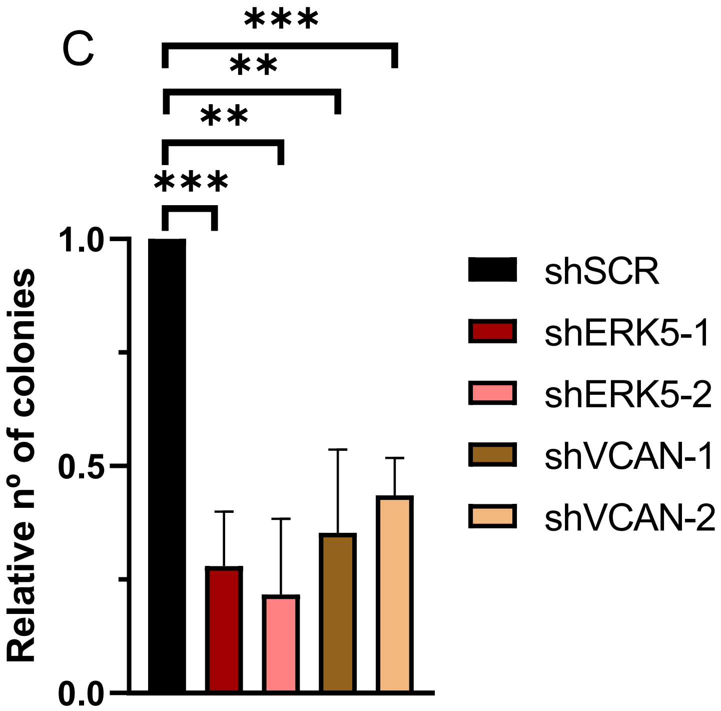

C
***
**
**
***
1.0
0.5
0.0
Relative nº of colonies
shSCR
shERK5-1
shERK5-2
shVCAN-1
shVCAN-2


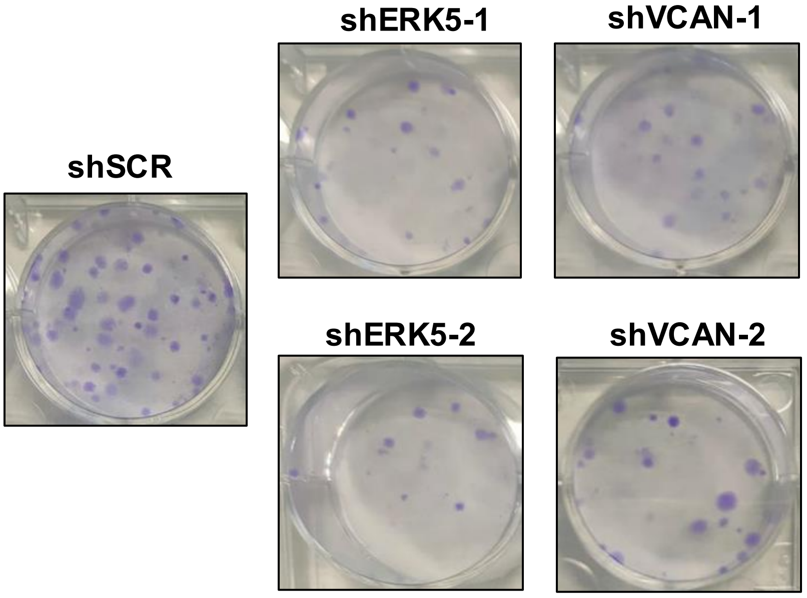

shERK5-1
shVCAN-1
shSCR
shERK5-2
shVCAN-2

**Supp. Fig. 4**

# EC

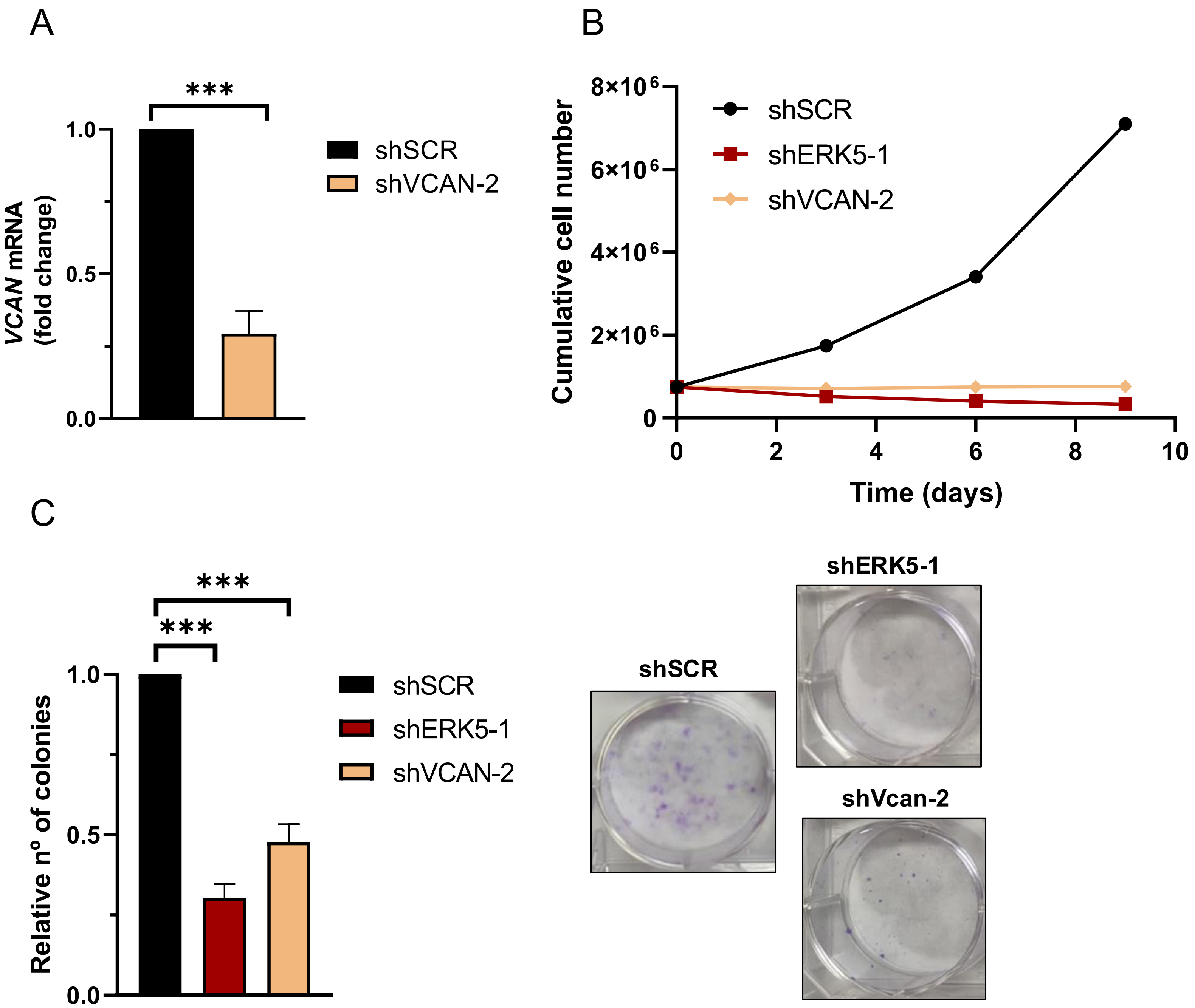
A
***
1.0
0.5
0.0
VCAN mRNA (fold change)
shSCR
shVCAN-2
B
8×10^6
6×10^6
4×10^6
2×10^6
0
Cumulative cell number
shSCR
shERK5-1
shVCAN-2
0
2
4
6
8
10
Time (days)
C
***
***
1.0
0.5
0.0
Relative nº of colonies
shSCR
shERK5-1
shVCAN-2
shSCR
shERK5-1
shVcan-2

Supp. Fig. 5

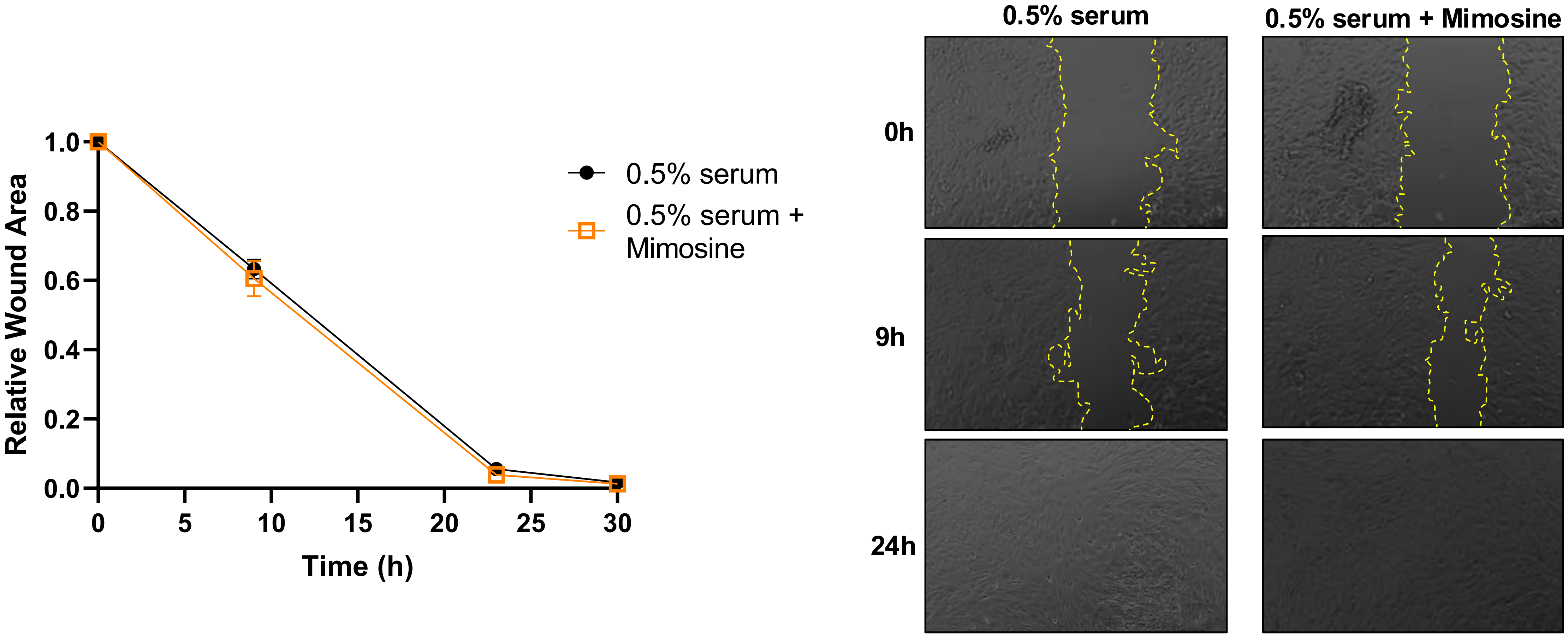
Relative Wound Area
1.0
0.8
0.6
0.4
0.2
0.0
0
5
10
15
20
25
30
Time (h)
0.5% serum
0.5% serum + Mimosine
0.5% serum
0.5% serum + Mimosine
0h
9h
24h

**Supp. Fig. 6**

A

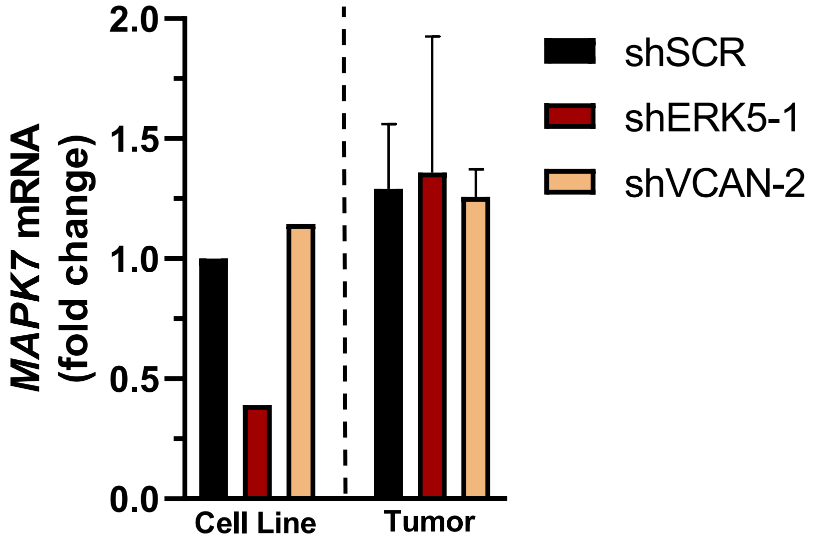

MAPK7 mRNA (fold change)
2.0
1.5
1.0
0.5
0.0
Cell Line
Tumor
shSCR
shERK5-1
shVCAN-2


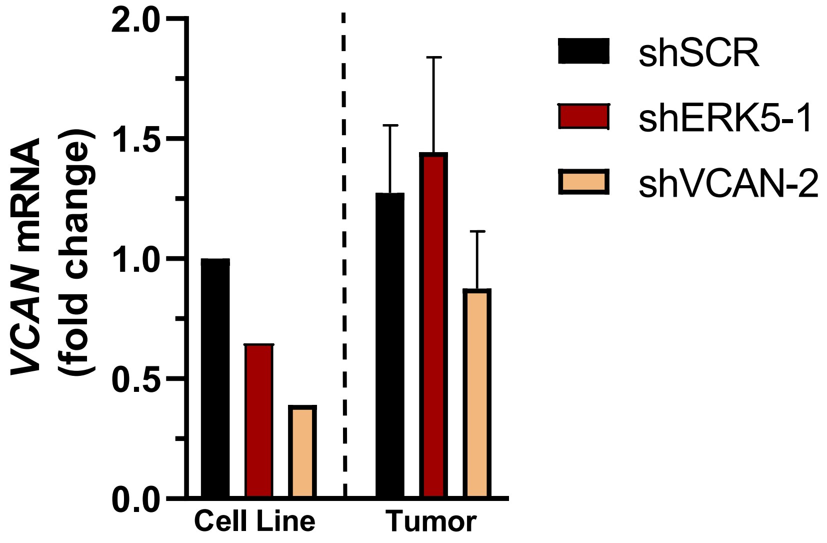

VCAN mRNA (fold change)
2.0
1.5
1.0
0.5
0.0
Cell Line
Tumor
shSCR
shERK5-1
shVCAN-2


B

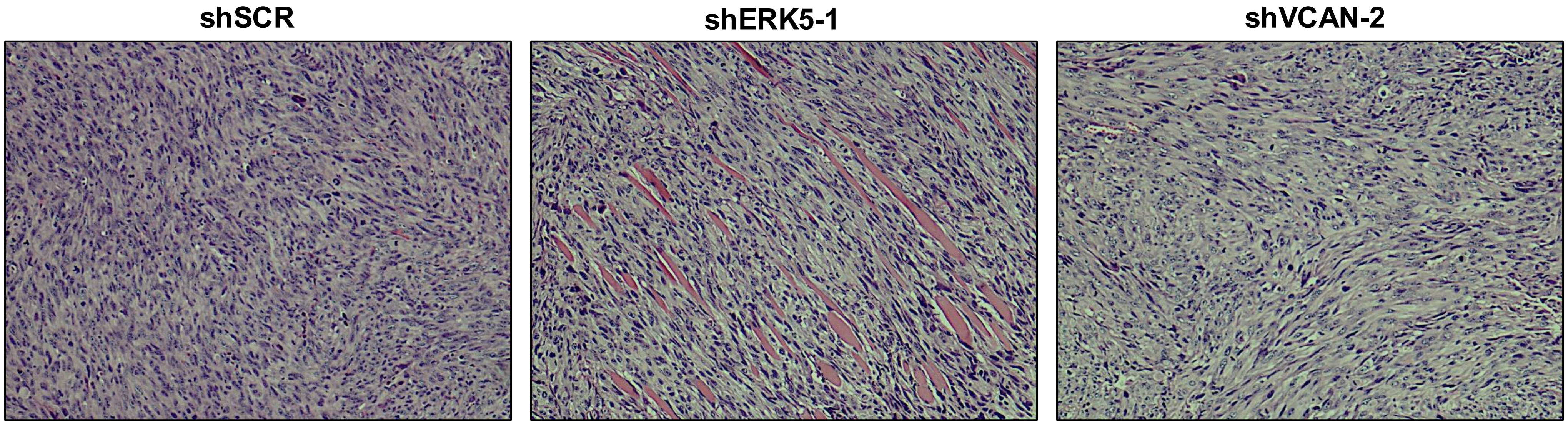

shSCR
shERK5-1
shVCAN-2

**Supp. Fig. 7**

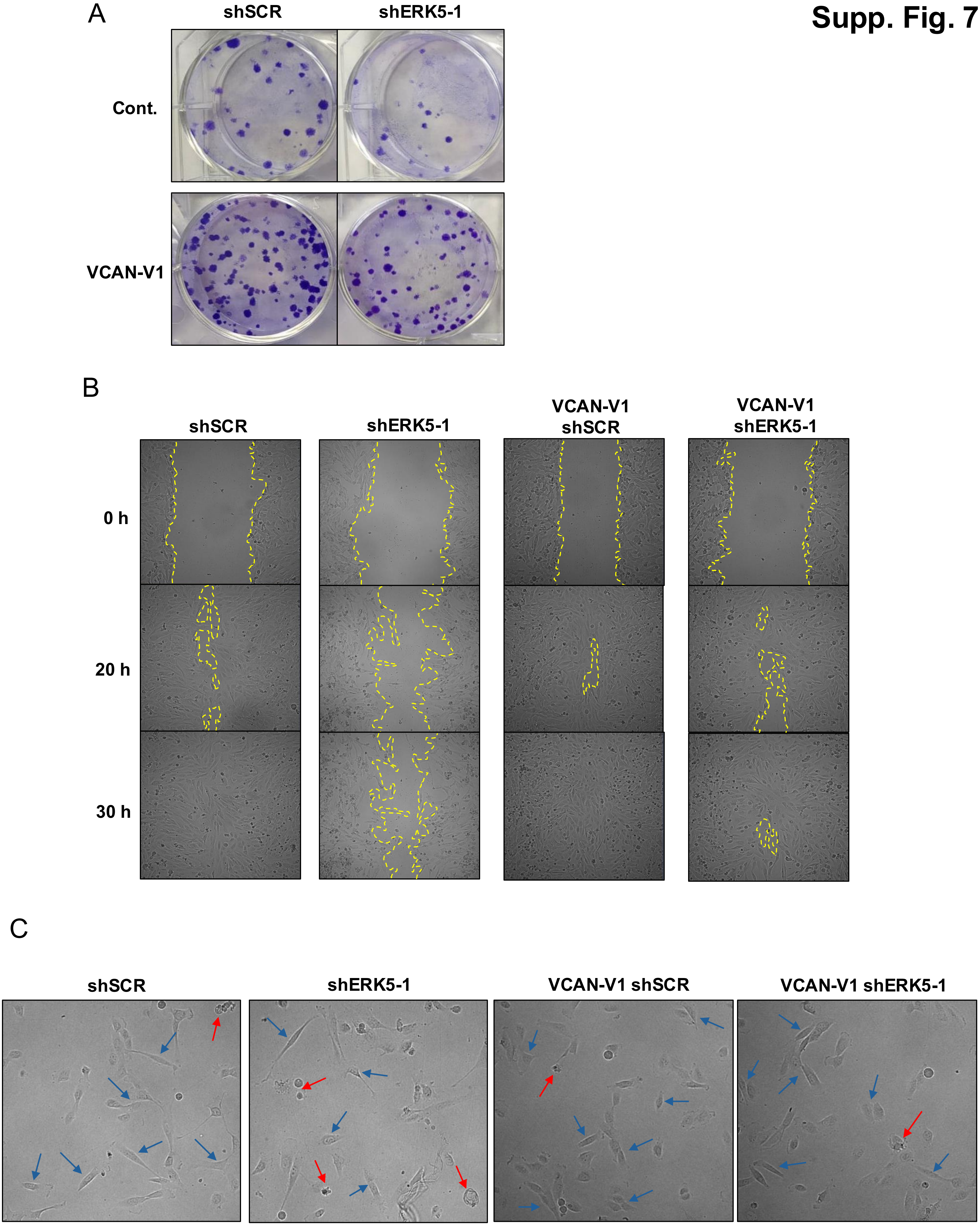
A
shSCR
shERK5-1
Cont.
VCAN-V1
B
shSCR
shERK5-1
VCAN-V1 shSCR
VCAN-V1 shERK5-1
0 h
20 h
30 h
C
shSCR
shERK5-1
VCAN-V1 shSCR
VCAN-V1 shERK5-1

Supp. Fig. 8

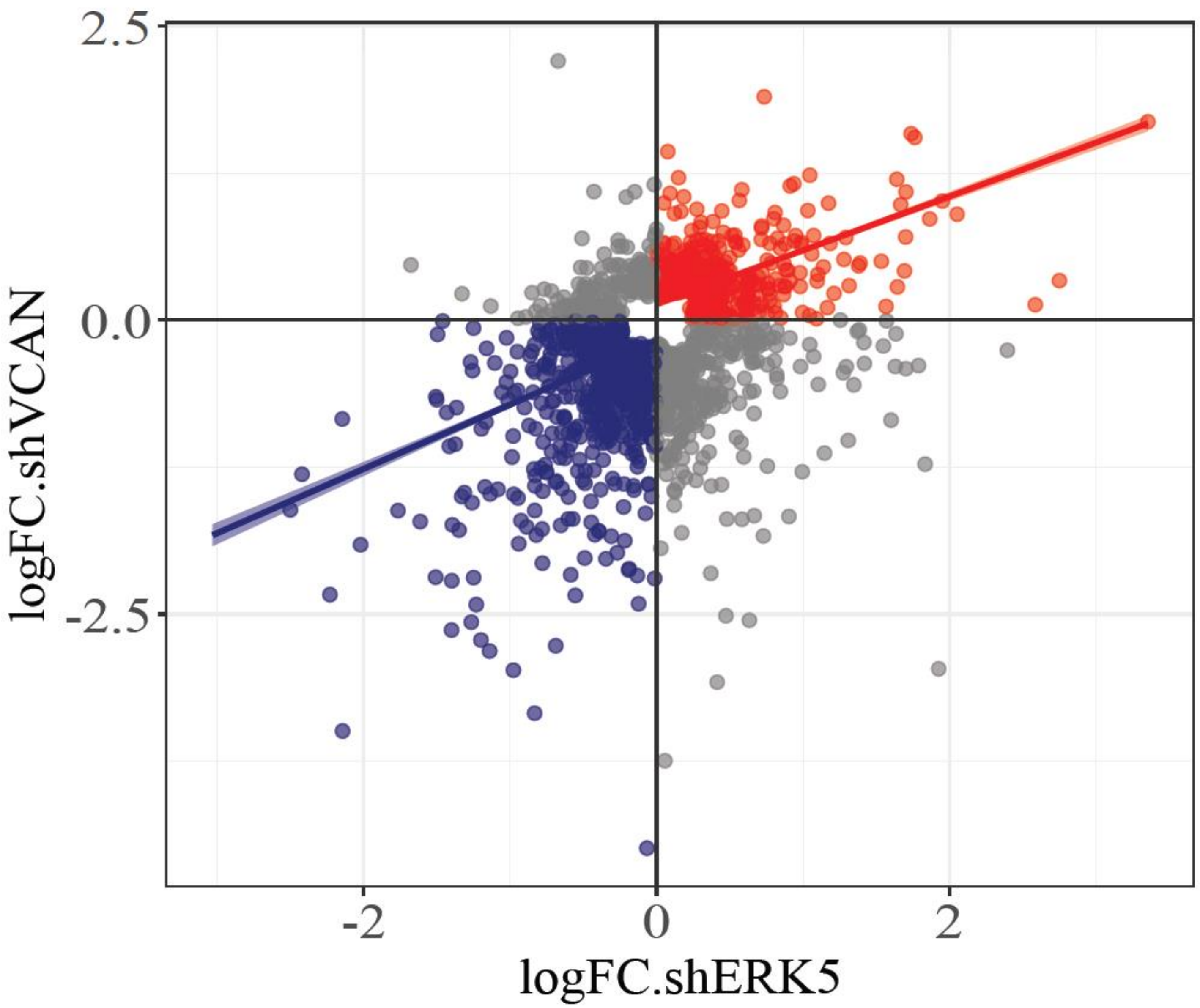

**Supp. Fig. 9**

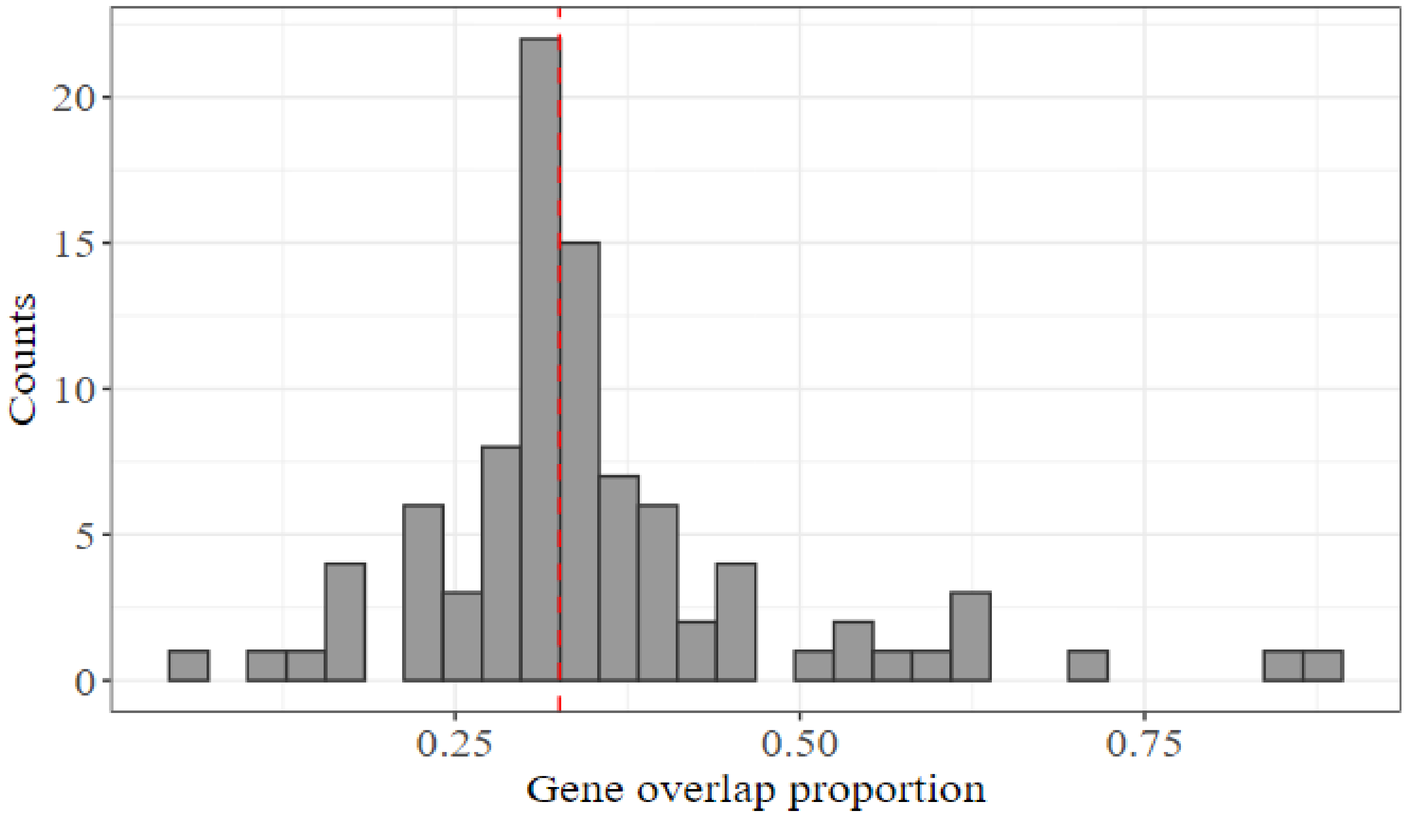

**Supp. Fig. 10**

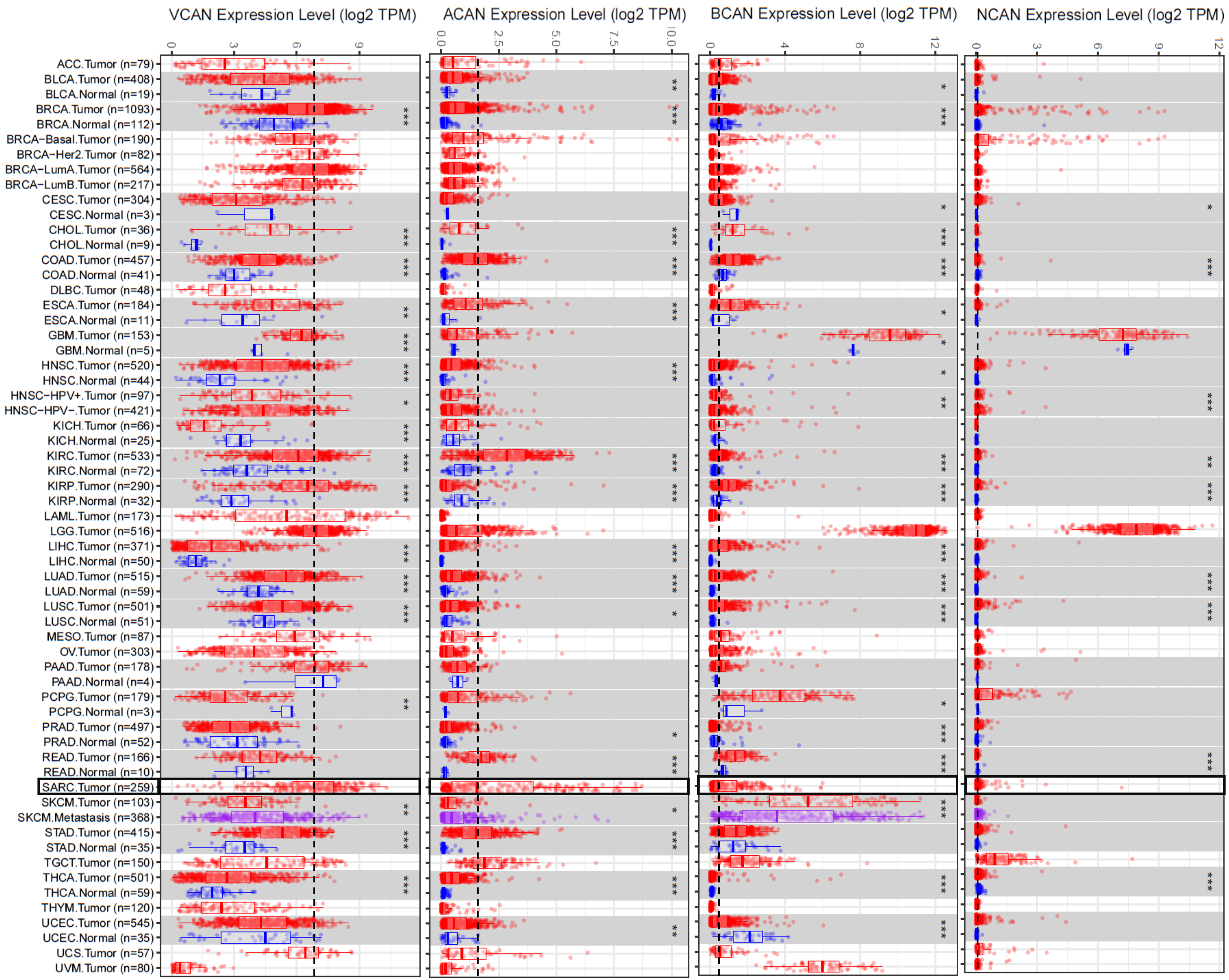
VCAN Expression Level (log2 TPM)
ACAN Expression Level (log2 TPM)
BCAN Expression Level (log2 TPM)
NCAN Expression Level (log2 TPM)
0 3 6 9
0.0 2.5 5.0 7.5 10.0
0 4 8 12
0 3 6 9 12
ACC.Tumor (n=79)
BLCA.Tumor (n=408)
BLCA.Normal (n=19)
BRCA.Tumor (n=1093)
BRCA.Normal (n=112)
BRCA−Basal.Tumor (n=190)
BRCA−Her2.Tumor (n=82)
BRCA−LumA.Tumor (n=564)
BRCA−LumB.Tumor (n=217)
CESC.Tumor (n=304)
CESC.Normal (n=3)
CHOL.Tumor (n=36)
CHOL.Normal (n=9)
COAD.Tumor (n=457)
COAD.Normal (n=41)
DLBC.Tumor (n=48)
ESCA.Tumor (n=184)
ESCA.Normal (n=11)
GBM.Tumor (n=153)
GBM.Normal (n=5)
HNSC.Tumor (n=520)
HNSC.Normal (n=44)
HNSC−HPV+.Tumor (n=97)
HNSC−HPV−.Tumor (n=421)
KICH.Tumor (n=66)
KICH.Normal (n=25)
KIRC.Tumor (n=533)
KIRC.Normal (n=72)
KIRP.Tumor (n=290)
KIRP.Normal (n=32)
LAML.Tumor (n=173)
LGG.Tumor (n=516)
LIHC.Tumor (n=371)
LIHC.Normal (n=50)
LUAD.Tumor (n=515)
LUAD.Normal (n=59)
LUSC.Tumor (n=501)
LUSC.Normal (n=51)
MESO.Tumor (n=87)
OV.Tumor (n=303)
PAAD.Tumor (n=178)
PAAD.Normal (n=4)
PCPG.Tumor (n=179)
PCPG.Normal (n=3)
PRAD.Tumor (n=497)
PRAD.Normal (n=52)
READ.Tumor (n=166)
READ.Normal (n=10)
SARC.Tumor (n=259)
SKCM.Tumor (n=103)
SKCM.Metastasis (n=368)
STAD.Tumor (n=415)
STAD.Normal (n=35)
TGCT.Tumor (n=150)
THCA.Tumor (n=501)
THCA.Normal (n=59)
THYM.Tumor (n=120)
UCEC.Tumor (n=545)
UCEC.Normal (n=35)
UCS.Tumor (n=57)
UVM.Tumor (n=80)

**Supp. 11**

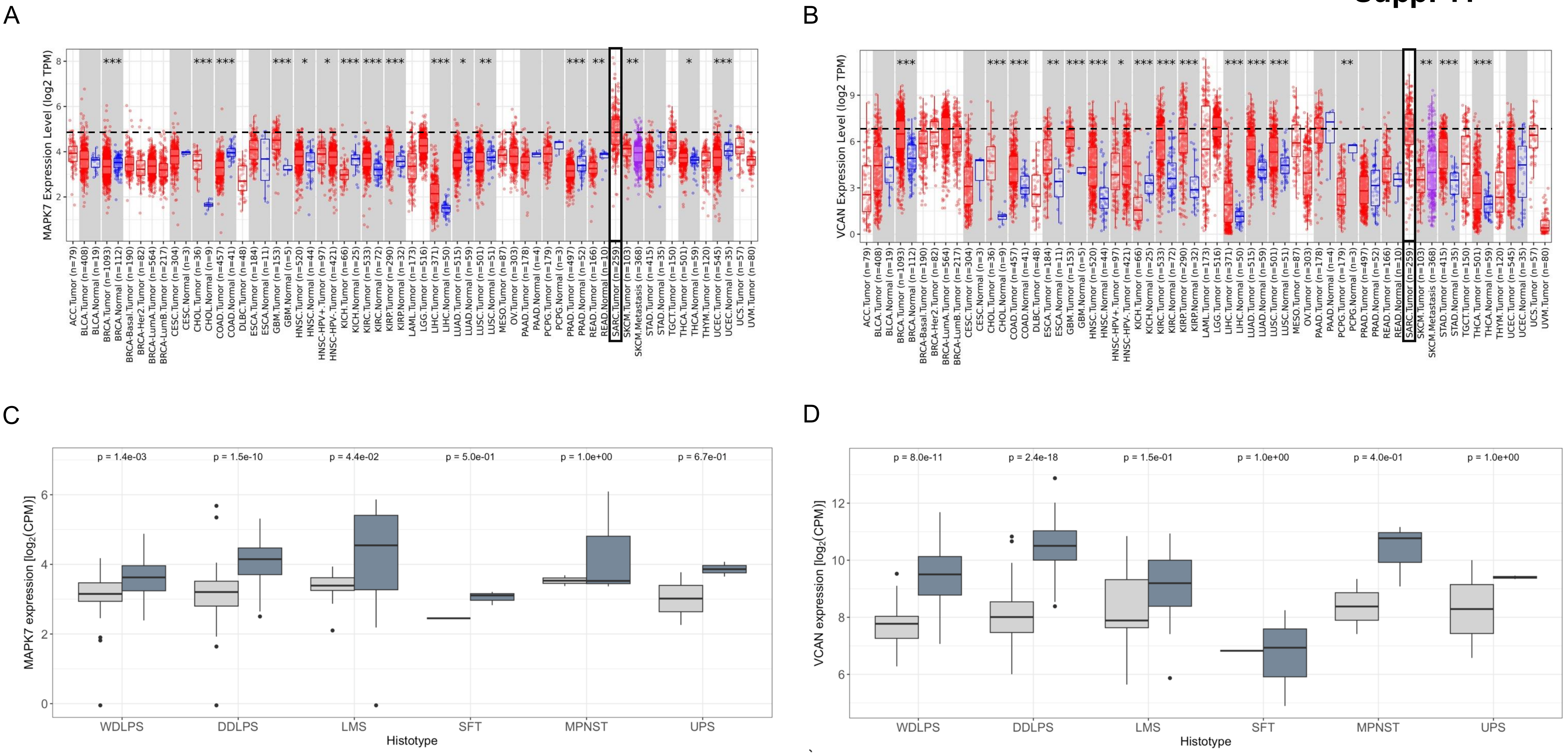
A
MAPK7 Expression Level (log2 TPM)
B
VCAN Expression Level (log2 TPM)
ACC.Tumor (n=79)
BLCA.Tumor (n=408)
BLCA.Normal (n=19)
BRCA.Tumor (n=1093)
BRCA.Normal (n=112)
BRCA-Basal.Tumor (n=190)
BRCA-Her2.Tumor (n=82)
BRCA-LumA.Tumor (n=564)
BRCA-LumB.Tumor (n=217)
CESC.Tumor (n=304)
CESC.Normal (n=3)
CHOL.Tumor (n=36)
CHOL.Normal (n=9)
COAD.Tumor (n=457)
COAD.Normal (n=41)
DLBC.Tumor (n=48)
ESCA.Tumor (n=184)
ESCA.Normal (n=11)
GBM.Tumor (n=153)
GBM.Normal (n=5)
HNSC.Tumor (n=520)
HNSC.Normal (n=44)
HNSC-HPV+.Tumor (n=97)
HNSC-HPV-.Tumor (n=421)
KICH.Tumor (n=66)
KICH.Normal (n=25)
KIRC.Tumor (n=533)
KIRC.Normal (n=72)
KIRP.Tumor (n=290)
KIRP.Normal (n=32)
LAML.Tumor (n=173)
LGG.Tumor (n=516)
LIHC.Tumor (n=371)
LIHC.Normal (n=50)
LUAD.Tumor (n=515)
LUAD.Normal (n=59)
LUSC.Tumor (n=501)
LUSC.Normal (n=51)
MESO.Tumor (n=87)
OV.Tumor (n=303)
PAAD.Tumor (n=178)
PAAD.Normal (n=4)
PCPG.Tumor (n=179)
PCPG.Normal (n=3)
PRAD.Tumor (n=497)
PRAD.Normal (n=52)
READ.Tumor (n=166)
READ.Normal (n=10)
SARC.Tumor (n=259)
SKCM.Tumor (n=103)
SKCM.Metastasis (n=368)
STAD.Tumor (n=415)
STAD.Normal (n=35)
TGCT.Tumor (n=150)
THCA.Tumor (n=501)
THCA.Normal (n=59)
THYM.Tumor (n=120)
UCEC.Tumor (n=545)
UCEC.Normal (n=35)
UCS.Tumor (n=57)
UVM.Tumor (n=80)
C
MAPK7 expression [log2(CPM)]
p = 1.4e-03
p = 1.5e-10
p = 4.4e-02
p = 5.0e-01
p = 1.0e+00
p = 6.7e-01
D
VCAN expression [log2(CPM)]
p = 8.0e-11
p = 2.4e-18
p = 1.5e-01
p = 1.0e+00
p = 4.0e-01
p = 1.0e+00
WDLPS
DDLPS
LMS
SFT
MPNST
UPS
Histotype

**Supplementary Table 1. List of antibodies used.**

| Antibody | Dilution | Manufacture | Reference | Use |
|---|---|---|---|---|
| ERK5 | 1:1000 | Cell Signaling Technology | 3372S | Western blot |
| VCAN | 1:5000 | Abcam | Ab19345 | Western blot |
| MEK5 | 1:1000 | Enzo Life Science | ADI-KAP-MA003-E | Western blot |
| HA | 1:1000 | Sigma-Aldrich | H9658 | Western blot |
| Vinculin | 1:3000 | Sigma-Aldrich | V9264 | Western blot |
| Nidogen 1 | 1:3000 | R&D Systems | AF2570 | Western blot |
| Calnexin | 1:1000 | Stressgen Bioreagents | SPC-127 | Western blot |
| Anti-mouse | 1:3000 | Cell Signaling Technology | 7076S | Western blot |
| Anti-rabbit | 1:3000 | Cell Signaling Technology | 7074S | Western blot |
| ERK5 | 1:500 | Abcam | ab196609 | Immunohistochemistry |
| VCAN | 1:2000 | Abcam | ab270445 | Immunohistochemistry |

**Supplementary Table 2. List of primers used.**

| Primer | Forward | Reverse |
|---|---|---|
| human *MAPK7* | AGCACTTTAAACACGACAAC | TAGACAGATTTGAATTCGCC |
| human *VCAN* | AGGTGGTCTACTTGGGGTGA | TGGTTGTAGCCTCTTTAGGTTT |
| human *CDKN1A* | ACTCTCAGGGTCGAAAACGG | CTTCCTGTGGGCGGATTAGG |
| human *GAPDH* | TCGTGGAAGGACTCATGACCA | CAGTCTTCTGGGTGGCAGTGA |
| mouse *Mapk7* | AGATCTGTCTATGTGGTACTG | CTGGTACAGGAAGTATCTCAC |
| mouse *Vcan* | AGGCGTCTACCGATGTGATG | CAGCGGCAAAGTTCAGAGTG |
| mouse *B2m* | GGTCTTTCTGGTGCTTGTCTCA | GTTCGGCTTCCCATTCTCC |

**Supplementary Table 3. Clinical and pathological characteristics of 222 patients with either primary retroperitoneal sarcoma (RPS) or primary extremity soft tissue sarcomas (ESTS) who were included in the retrospective cohort of the SARCOMCS clinical trial and were profiled with RNA-sequencing**. WDLPS: Well-differentiated liposarcoma; DDLPS: dedifferentiated liposarcoma; PLPS: pleomorphic liposarcoma; MLPS: myxoid liposarcoma; LMS: leiomyosarcoma; MPNST: malignant peripheral nerve sheath tumor; UPS: undifferentiated pleomorphic sarcoma; SS: synovial sarcoma; AS: angiosarcoma; SFT: solitary fibrous tumor; MFS: myxofibrosarcoma.

| **Site** | RPS | ESTS | All |
|---|---|---|---|
| **Sex** | | | |
| Female | 46 (34%) | 38 (44%) | 84 (38%) |
| Male | 89 (66%) | 49 (56%) | 138 (62%) |
| **Age (median, IQR)** | 63 (54.5-72) | 59 (46-71.5) | 62 (50-72) |
| **Histology** | | | |
| WDLPS | 40 (30%) | 0 | 40 (18%) |
| DDLPS | 69 (51%) | 4 (5%) | 73 (33%) |
| PLPS | 0 | 5 (6%) | 5 (2%) |
| MLPS | 0 | 14 (16%) | 14 (6%) |
| LMS | 17 (13%) | 7 (8%) | 24 (11%) |
| MPNST | 3 (2%) | 2 (2%) | 5 (2%) |
| UPS | 2 (1%) | 23 (26%) | 25 (11%) |
| SS | 0 | 3 (3%) | 3 (2%) |
| AS | 0 | 1 (2%) | 1 (1%) |
| SFT | 4 (3%) | 7 (8%) | 11 (5%) |
| MFS | 0 | 21 (24%) | 21 (9%) |
| **Grading** | | | |
| I | 44 (33%) | 21 (24%) | 65 (29%) |
| II | 57 (42%) | 20 (23%) | 77 (35%) |
| III | 34 (25%) | 46 (53%) | 80 (36%) |
| **Neoadjuvant treatment** | | | |
| Chemotherapy | 7 (5%) | 15 (17%) | 22 (9%) |
| Radiotherapy | 15 (11%) | 9 (10%) | 24 (11%) |

**Supplementary Table 4. Clinicopathological data from Complejo Hospitalario Universitario de Albacete sarcoma patients cohort.**

| ID | Age | Sex | Diagnosis | Location | ERK5 Intensity | VCAN Intensity |
|---|---|---|---|---|---|---|
| **Patient 01** | 77 | M | Leiomyosarcoma | Soft parts of the lower extremity | +++ | +++ |
| **Patient 02** | 68 | F | Leiomyosarcoma | Uterus | +++ | ++ |
| **Patient 03** | 64 | F | Leiomyosarcoma | Mediastinum | 0 | 0 |
| **Patient 04** | 44 | M | Leiomyosarcoma | Rectum | + | 0 |
| **Patient 05** | 58 | M | Leiomyosarcoma | Axilla | 0 | 0 |
| **Patient 06** | 48 | F | Leiomyosarcoma | Uterus | 0 | 0 |
| **Patient 07** | 49 | F | Leiomyosarcoma | Lung | + | + |
| **Patient 08** | 35 | F | Leiomyosarcoma | Uterus | + | +++ |
| **Patient 09** | 58 | F | Leiomyosarcoma | Uterus | + | 0 |
| **Patient 10** | 56 | F | Leiomyosarcoma | Uterus | + | 0 |
| **Patient 11** | 55 | M | Undifferentiated pleomorphic sarcom | Soft parts of the chest wall | ++ | ++ |
| **Patient 12** | 79 | F | Undifferentiated pleomorphic sarcom | Soft parts of the lower extremity | ++ | ++ |
| **Patient 13** | 72 | F | Undifferentiated pleomorphic sarcom | Meninges | 0 | 0 |
| **Patient 14** | 71 | M | Undifferentiated pleomorphic sarcom | Pelvis | + | + |
| **Patient 15** | 70 | M | Undifferentiated pleomorphic sarcom | Soft parts of the upper extremity | ++ | 0 |
| **Patient 16** | 70 | M | Undifferentiated pleomorphic sarcom | Rectum | ++ | ++ |
| **Patient 17** | 48 | F | Undifferentiated pleomorphic sarcom | Soft parts of the lower extremity | +++ | +++ |
| **Patient 18** | 76 | M | Undifferentiated pleomorphic sarcom | Soft parts of the lower extremity | +++ | ++ |
| **Patient 19** | 78 | M | Undifferentiated pleomorphic sarcom | Soft parts of the neck region | ++ | + |
